# Loss of TAFAZZIN leads to perturbation of amino acid metabolism and reduction of collagen synthesis

**DOI:** 10.64898/2026.07.19.739395

**Authors:** Abu Ramim, Tyler Ralph-Epps, Linh Vo, Hyejeong Jang, Janaka S.S. Liyanage, Kaitlin Lowran, Miriam L. Greenberg

## Abstract

Barth syndrome is a life-threatening genetic disorder caused by mutations in the TAFAZZIN (TAZ) gene, which disrupt remodeling of cardiolipin in mitochondria. The disease is associated with cardiac and skeletal myopathy, neutropenia, fatigue, and metabolic dysfunction. Previous studies showed that loss of TAZ decreases pyruvate dehydrogenase activity, reduces glucose flux into the TCA cycle, and impairs fatty acid metabolism. To test the hypothesis that amino acid (AA) metabolism may be altered to compensate for these deficiencies, we characterized AA metabolism in TAZ-deficient mouse myoblasts (TAZ-KO). Levels of branched-chain amino acids (BCAAs) were reduced, while proline levels were increased in TAZ-KO cells. Levels of proline dehydrogenase and glutamate dehydrogenase, which convert proline to TCA cycle intermediates, were increased. ^13^C_5_-proline isotope tracing demonstrated elevated conversion of proline into glutamate and TCA cycle intermediates. SILAC analysis using [U-^13^C_6_, ^15^N_2_]-Lys and [U-^13^C_6_]-Arg revealed decreased synthesis of collagen and proteins associated with extracellular matrix (ECM). Gene expression and protein analyses revealed reduced collagen expression, lower total collagen content, decreased collagen crosslinking enzymes, decreased proline hydroxylation and reduced synthesis of new collagen and cell-adhesion proteins. SILAC analysis using [U-^13^C_6_, ^15^N_2_]-proline also showed diminished incorporation of proline into newly synthesized ECM proteins. Together, our findings reveal that loss of TAZ leads to increased proline catabolism to the TCA cycle, decreased incorporation of proline into collagen, and impaired collagen synthesis and ECM remodeling.

## 1. Introduction

Barth syndrome (BTHS) is a rare X-linked genetic disorder caused by mutations in the TAFAZZIN (TAZ) gene, which encodes a mitochondrial transacylase that remodels cardiolipin (CL) [1–3]. The disease is characterized by cardiomyopathy, skeletal myopathy, neutropenia, fatigue and exercise intolerance, and metabolic abnormalities. The hallmark biochemical defect is the accumulation of monolyso-CL (MLCL) and near absence of unsaturated CL species [2–6]. The precise connection between TAZ deficiency and the pathology in BTHS is still unknown, underscoring the importance of clarifying how loss of TAZ contributes to metabolic dysfunction in the disorder.

We previously reported that pyruvate dehydrogenase (PDH)—the gatekeeper enzyme routing pyruvate into the TCA cycle—is impaired in TAZ-deficient C2C12 myoblasts (TAZ-KO) and cardiac tissue [7–9]. PDH deficiency was also reported in the TAZ-deficient BTHS mouse model, confirming PDH dysfunction as a consistent feature of BTHS [7, 9]. PDH activity is controlled by reversible phosphorylation—PDH kinase (PDK) inactivates the enzyme by phosphorylation, and PDH phosphatase (PDP) reactivates it via dephosphorylation. The observed reduction in PDP1 activity and elevation in PDK4 expression underlie the mechanism whereby loss of TAZ impairs PDH regulation [8]. Consistent with decreased PDH activity, glucose flux to the TCA cycle is reduced [7]. Heart and skeletal muscle rely on fatty acid oxidation (FAO) as a major fuel pathway; when glucose metabolism is compromised, these tissues rely even more on FAO to meet their energy demands [10]. In TAZ-KO fibroblasts, the *TAZ* mutant mouse heart, and patient-derived induced pluripotent stem cell (iPSC) cardiomyocytes, expression of FAO-associated genes and proteins is reduced [11]. Furthermore, FA supplementation did not restore respiration [11]. These results suggest that both glucose and FA metabolism are impaired in TAZ-deficient cells, contributing to the cardiac and skeletal muscle dysfunction characteristic of BTHS.

As amino acid (AA) metabolism is an alternate route to generate energy via the TCA cycle, we considered the possibility that AA metabolism may be altered to compensate for the deficiencies in glucose and FA metabolism. To this end, we examined steady-state AA levels in TAZ-KO cells. LC–MS profiling revealed decreased levels of branched chain AAs (BCAAs) leucine, isoleucine, and valine and increased proline in TAZ-KO cells. BCAAs are of particular interest, as they account for ∼35% of essential AAs in muscle proteins and are primarily catabolized in skeletal muscle, while most other AAs are metabolized in the liver [12–14]. BCAAs are catabolized to generate acetyl-CoA and succinyl-CoA, connecting BCAAs to mitochondrial energy production and anabolic signaling [15]. Proline, a nonessential AA (NEAA), can be metabolized to glutamate and subsequently to α-ketoglutarate to replenish the TCA cycle; alternatively, proline can be directed toward collagen production and redox regulation based on cellular needs [16]. Increased flux from glutamine to proline has been reported in BTHS [17].

In addition to its anaplerotic role in replenishing the TCA cycle, proline is also crucial for collagen synthesis and muscle development. Muscle differentiation depends on proper support from the ECM, where remodeling maintains cell adhesion, signaling, and structural organization [18]. Collagen, the most abundant ECM protein, provides the scaffold for tissue integrity and force transmission during myotube formation [19, 20]. Proline constitutes about 13% of the AA content of collagen, and impaired proline metabolism or collagen synthesis can disrupt ECM remodeling, leading to weakened cell adhesion, incomplete differentiation, and defective fiber maturation [21, 22]. This could be linked to reduced muscle strength, impaired regeneration, and skeletal myopathy over time which is a major clinical manifestation of BTHS [5, 18, 23–25].

The current study was carried out to determine how TAZ deficiency perturbs AA metabolism. To this end, we analyzed the expression of key BCAA catabolic enzymes and traced flux of [U-^13^C]-BCAAs into TCA cycle intermediates to determine if decreased levels of BCAAs reflected increased utilization to replenish TCA cycle intermediates. Further, we performed [U-^13^C₅]-proline tracing and measured the expression and activity of enzymes to test for a bottleneck in proline entry into the TCA cycle. Finally, we assessed collagen synthesis and incorporation of [U-^13^C_5_]-proline into newly synthesized collagen and ECM proteins. Our findings indicate that TAZ deficient cells exhibit: (1) increased proline catabolism, characterized by elevated expression of proline catabolic enzymes and increased incorporation of proline into the TCA cycle; and (2) decreased collagen content, proline hydroxylation, reduced expression of collagen crosslinking enzymes, and diminished incorporation of proline into collagen and ECM remodeling proteins.

## 2. Methods

### 2.1. Cell line and growth conditions

Wild type (WT) C2C12 mouse myoblasts (serve as the control group) and TAZ-KO knockout C2C12 mouse myoblasts were generated previously as described [3, 7]. Growth medium consisted of DMEM (Gibco) containing 10% fetal bovine serum (FBS; Optima), 2 mM glutamine (Gibco), penicillin (100 units/mL), and streptomycin (100 μg/mL) (Invitrogen). Cells were grown at 37°C in a humidified incubator with 5% CO_2_.

### 2.2. RNA extraction and qPCR

cDNA was generated from total RNA using a SuperScript™ kit (Invitrogen). Quantitative rt-PCR was performed on a QuantStudio 3 system (Applied Biosystems) with PowerUp™ SYBR™ green reagent (Thermo Fisher). Relative mRNA expression levels were calculated using the 2–^ΔΔ^CT method. *β*-actin was used as the internal reference gene. The primer sequences used are provided in Table 1.

**Table 1:**
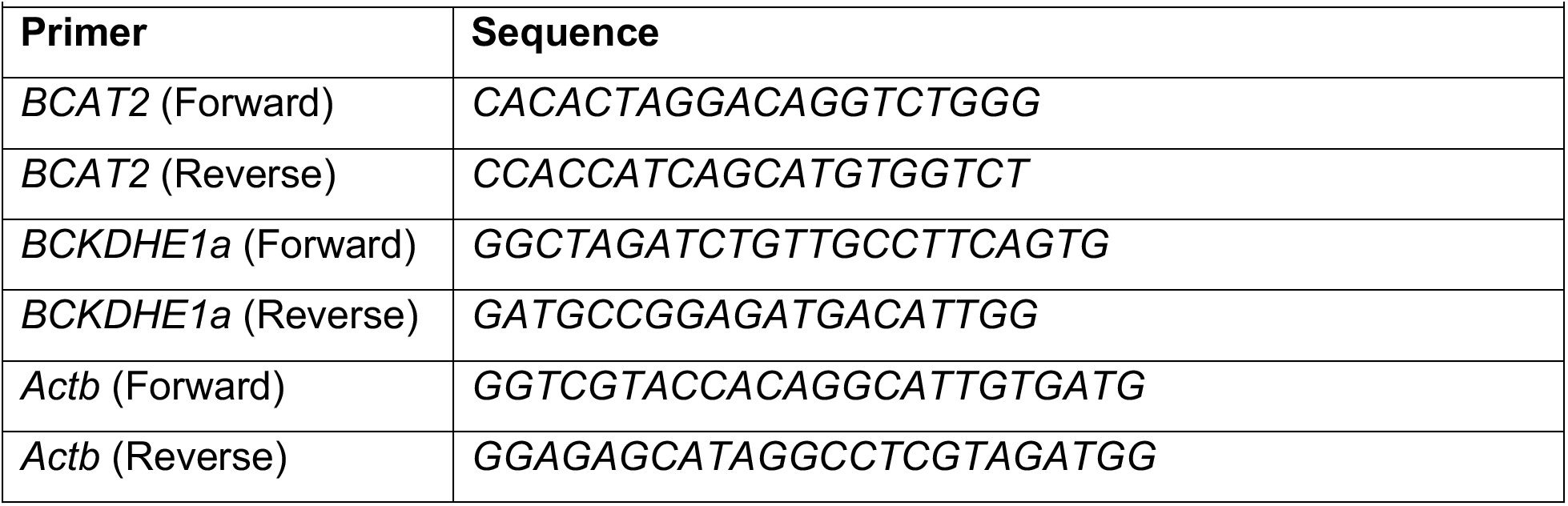
List of qPCR primers used.

### 2.3. Western blot

Proteins were extracted using RIPA buffer supplemented with protease and phosphatase inhibitor cocktail (ChemCruz). A total of 20-50 μg of denatured protein from each sample were loaded onto 8–12% SDS-PAGE gels and separated by electrophoresis. For each condition, at least three or four biological replicates were used. Proteins were then transferred to PVDF membranes by BIO-RAD heat transfer. Transferring the Col1a1 protein (220 kDa) required an overnight, low-voltage Western blot procedure at a temperature of 4°C. Total protein was quantified from each sample using a No-Stain labeling reagent (Invitrogen) and then used for total protein normalization (TPN). The membranes were subsequently incubated with primary antibodies against BCAT2 (1:1000, 16417-1-AP; Proteintech), BCKDHE1a (1:1000, 90198; Cell Signaling), p-BCKDHE1a (1:1000, 40368; Cell Signaling), SLC25A44 (1:1000, LS-C750336-20, LS BIO), LAT1 (D10) (1:1000, sc-374232, Santa Cruz Biotechnology), PRODH (1:2000, 22980-1-AP, Proteintech), GDH (1:1000, D9F7P; Cell Signaling), LOX (1:1000, 58135; Cell signaling technology), SLC6A20 (1:1000, 30843-1-AP; Proteintech), SNAT2 (1:1000, 25928-1-AP; Proteintech) and Col1a1 (1:1000, 72026; Cell signaling technology). Anti-mouse or anti-rabbit secondary antibodies (1:10,000) were incubated for 1 h at room temperature, and membranes were developed using SuperSignal™ West Pico PLUS or West Atto ECL (Thermo Scientific).

### 2.4. Collagen assay

Collagen content was assessed using the Hydroxyproline Assay Kit (MAK463, Sigma-Aldrich). Hydroxyproline is a modified AA found almost exclusively in collagen, constituting ∼13% of its total weight, making it a reliable marker of collagen levels. In this assay, hydroxyproline is first oxidized to a pyrrole intermediate, which then reacts with a dye reagent to form a pink product measurable at 560 nm (Spectramax i3x, Molecular Devices). The kit provides a linear detection range of 0.5–50 μg/mL hydroxyproline in a 96-well plate format. WT and TAZ-KO cells (N = 4 per group, with 3 technical replicates each) were analyzed, and results were normalized to protein concentration.

### 2.5. Isotope tracing

#### 2.5.1. Media preparation

To trace the incorporation of BCAA into the TCA cycle using LC-MS, DMEM, ATCC^®^30-2002 lacking BCAAs was prepared and supplemented with isotope labeled BCAAs along with standard FBS: 0.8mM (0.105 g/L) ^13^C_6_-leucine (CLM-2262-H-PK, Cambridge Isotope), 0.8mM (0.105 g/L) ^13^C_6_-isoleucine (CLM-2248-H-PK, Cambridge Isotope), 0.8mM (0.094 g/L) ^13^C₅-valine, and (CLM 2249-H-PK, Cambridge Isotope). For the glucose flux assay, glucose-free media was supplemented with 25 mM ^13^C_6_-glucose (CLM-1396-1). For control conditions, unlabeled AAs or glucose were added instead with similar concentration. For the proline flux assay, 2mM [U-^13^C₅]-proline (CLM-2260-H-PK) was added.

#### 2.5.2. Growth condition

In 6-well plates, 10,000 cells/cm^2^ were seeded per well in DMEM containing unlabeled AAs or glucose, with four biological replicates (N=4). For BCAA and glucose flux assay, plates were incubated at 37°C with CO_2_ for 44 hours in unlabeled media, after which the media were replaced with DMEM containing ^13^C-labeled BCAAs or glucose. Blank and control wells received unlabeled media. Plates were further incubated for 4 hours for BCAA/glucose tracing. For the proline flux assay, the procedure and processing were similar except seeding density and longer labeling period: 20000 cells/cm2 seeded and cultured in unlabeled media for 24 hours, then switched to labeled media and incubated for an additional 8 hours.

#### 2.5.3. Sample preparation

After incubation in ^13^C-labeled media (BCAA/glucose/proline), 200 µl of medium were collected from each well, centrifuged at 1000 rpm for 5 minutes, and immediately snap frozen. All 6-well plates were then washed twice with ice-cold 500 µl of 0.9% NaCl and plates were snap frozen immediately in liquid nitrogen. Blank wells without cells were processed in parallel under same conditions. All samples were transported on dry ice to the Van Andel Institute Metabolomics Core for metabolomic flux analysis.

#### 2.5.4. Metabolite extraction: adherent cells

For metabolomics analysis, an acetonitrile-methanol-water (4:4:2 v/v) extraction was used such that final extraction solvent to sample ratios were based on a standardized number of cells per 1 mL of extraction solvent. Equal numbers of cells were used across biological replicates and conditions (WT and TAZ-KO). The proper volume of ice-cold 4:4:2 acetonitrile (A955, Fisher): methanol (A456, Fisher): water (W6, Fisher) was added to all wells of a 6-well plate chilled on wet ice. Wells were sequentially scraped, and the 4:4:2 solvent-cell mixture was transferred to an empty tube and kept on wet ice. After solvent addition, extracts were vortexed for 10 s, sonicated in a water bath for 5 min, and incubated on wet ice for 1 hour. Incubation was followed by a 10-minute centrifugation at 17,000 g and 4°C to collect the solid protein pellet at the bottom of the tube and separate it from the metabolite-containing supernatant. Next, 1.6 million cell equivalents were aliquoted from the supernatant, dried under vacuum, and resuspended in 50 uL LCMS-grade water (for ion-paired analysis) or 50 uL 1:1 water: acetonitrile (for metabolomics analysis). A mix of deuterated standards from Cambridge Isotope Laboratories (listed in the table 2) was added as an internal standard to all metabolomics samples at a final concentration of 1 ug/mL to monitor instrument performance across each injection.

**Table 2:**
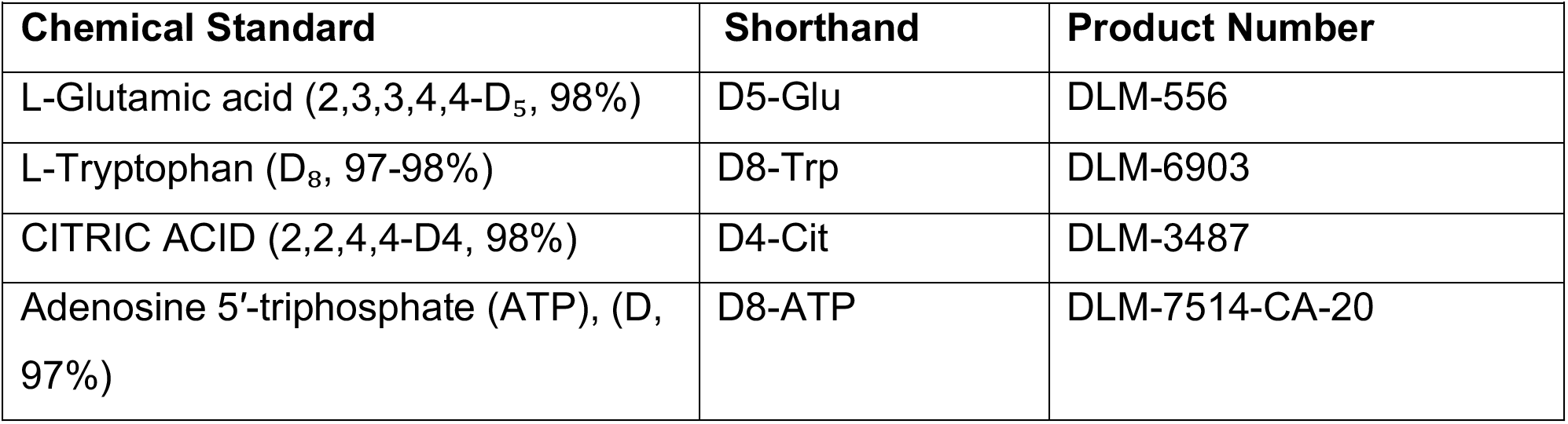
List of deuterated standards used as an internal standard.

| **Table 2: List of deuterated standards used as an internal standard** |  |  |
| --- | --- | --- |
| **Chemical Standard** | **Shorthand** | **Product Number** |
| L-Glutamic acid (2,3,3,4,4-D<sub>5</sub>, 98%) | D5-Glu | DLM-556 |
| L-Tryptophan (D<sub>8</sub>, 97-98%) | D8-Trp | DLM-6903 |
| CITRIC ACID (2,2,4,4-D<sub>4</sub>, 98%) | D4-Cit | DLM-3487 |
| Adenosine 5'-triphosphate (ATP), (D, 97%) | D8-ATP | DLM-7514-CA-20 |

#### 2.5.5. Metabolite extraction: media

For metabolomics analysis, an acetonitrile-methanol-water (4:4:2 v/v) extraction was used. Samples were homogenized in ice-cold 4:4:2 acetonitrile (A955, Fisher): methanol (A456, Fisher): water (W6, Fisher) such that final extraction solvent to sample ratios were 40 uL of sample to 1 mL of extraction solvent. After solvent addition, extracts were vortexed for 10 s, sonicated in a water bath for 5 min, and incubated on wet ice for 1 hour. Incubation was followed by a 10-minute centrifugation at 17,000 g and 4°C to collect the solid protein pellet at the bottom of the tube and separate it from the metabolite-containing supernatant. Next, 32 uL biofluid equivalents were aliquoted from the supernatant, dried under vacuum, and resuspended in 50 uL LCMS-grade water. A mix of deuterated standards from Cambridge Isotope Laboratories (listed in the table 2) was added as an internal standard to all metabolomics samples at a final concentration of 1 ug/mL to monitor instrument performance across each injection.

#### 2.5.4 Ion-paired Single Chromatography Profiling & Tracing

Samples were analyzed with a Vanquish liquid chromatography system coupled to an Orbitrap Exploris 240 (Thermo Fisher Scientific) using an H-ESI (heated electrospray ionization) source in negative mode. 2 μL of each standard and/or sample was injected and run through a 24-minute reversed-phase chromatography Acquity Premier HSS T3 Column (1.8 μm, 2.1mm × 150mm, 186009472, Waters, Eschborn, Germany) combined with a VanGuard cartridge (1.8 μm, 2.1 mm × 5 mm, 186009473, Waters). Mobile phase A consisted of LC/MS grade water (W6, Fisher) with 3% LC/MS grade methanol (A456, Fisher), mobile phase B was LC/MS grade methanol and both mobile phases contained 10 mM tributylamine (90780, Sigma), 15 mM LC/MS grade acetic acid (A11350, Fisher), and 0.01% medronic acid (v/v, 5191-4506, Agilent). For the wash gradient, mobile phase A was kept the same, and mobile phase B was 99% LC/MS grade acetonitrile (A955, Fisher). Column temperature was kept at 35 °C, flow rate was held at 0.25 mL/min, and the chromatography gradient was as follows: 0-2.5 min held at 0% B, 2.5-7.5 min from 0% B to 20% B, 7.5-13 min from 20% B to 45% B, 13-20 min from 45% B to 99% B, and 20-24 min held at 99% B. A 16 minute wash gradient was run in reverse flow direction between every injection to back-flush the column and to re-equilibrate solvent conditions as follows: 0-3 min held at 100% B and 0.25 mL/min, 3-3.5 min held at 100% B and ramp to 0.8 mL/min. 3.5-7.35 min held at 100% B and 0.8 mL/min, 7.35-7.5 held at 100% B and ramp to 0.6 mL/min, 7.5-8.25 from 100% B to 0% B and ramp to 0.4 mL/min, 8.25-15.5 min held at 0% B and ramp to 0.25 mL/min, and 15.5-16 min held at 0% B and 0.25 mL/min. Mass spectrometer parameters were: source voltage −2500V, sheath gas 60, aux gas 19, sweep gas 1, ion transfer tube temperature 320°C, and vaporizer temperature 250°C. Full scan data were collected using the orbitrap with a scan range of 70-850 m/z at a resolution of 240,000 and RF lens at 35%. Fragmentation was induced in the orbitrap using assisted higher-energy collisional dissociation (HCD) collision energies at 15, 30, and 45%. Orbitrap resolution was 15,000, the isolation window was 2 m/z, and data dependent scans were capped at 5 scans. Targeted mass MS2 triggers were included for a panel of compounds, displayed in Supporting Table s1.

### 2.6. Stable-isotope labeling (SILAC) to assess protein synthesis

#### 2.6.1. Media preparation

For global protein synthesis, DMEM lacking lysine (Lys) and arginine (Arg) was supplemented with isotope-labeled AAs together with dialyzed FBS (A5670401, Gibco): 146 mg/L [U-^13^C_6_, ^15^N_2_]-Lys (88209, Thermo Scientific) and 84 mg/L [U-^13^C_6_]-Arg (88210, Thermo Scientific). For collagen and ECM analysis, media was supplemented with 2mM ^13^C₅-proline (CLM-2260-H-PK, Cambridge Isotope Inc).

#### 2.6.2. Growth condition

WT and TAZ-KO cells (N = 4 per group) were seeded in “light” DMEM containing unlabeled media at a density of 10,000 cells/cm² for 12 hours (8,000 cells/cm² for 15 hours for proline flux to collagen/ECM remodeling protein) in 100 mm plates. After this period, the medium was switched to “heavy” medium containing labeled isotopes ([U-^13^C_6_, ^15^N_2_]-Lys and [U-^13^C_6_]-Arg for global protein synthesis, or ^13^C₅-Pro for collagen/ECM protein synthesis). Cells were cultured in heavy media for ∼24 hours for global protein synthesis and ∼80 hours for proline incorporation into collagen and ECM proteins.

#### 2.6.3. Sample preparation

At harvest, cells were washed twice with cold PBS, lysed in 500 µl of 8 M urea buffer with protease inhibitors (8 M urea, 50 mM Tris-HCl, pH 8.5), scraped, and collected into 1.5 ml Eppendorf tubes. Samples were transported on ice to the Karmanos Cancer Institute Proteomics Core Facility for mass spectrometry analysis. Samples were buffered with 20 mM triethylammonium bicarbonate (TEAB, Honeywell Fluka cat# 60-044-974) then reduced with 5 mM DL-dithiothreitol (DTT, Sigma cat# D5545) for 30 mins at 37°C, alkylated with 15 mM iodoacetamide (IAA, Sigma cat# I1149) for 30 mins at room temperature in the dark, then 5mM DTT was added to stop alkylation. Proteins in the samples were precipitated using a solution of 90% methanol (MeOH) and 100 mM TEAB. Precipitates were washed with 80% MeOH and 10 mM TEAB and then digested with 0.5 µg of trypsin (Promega, V5113) for 2 hours at 47°C. Digested samples were then dried in a speed vacuum system and dissolved in 50uL of 0.1% Formic acid (FA, ThermoScientific cat#T85170).

#### 2.6.4. Mass-spectrometry conditions

Mass spectrometry was performed using a Thermo Easy-nLC 1200 UHPLC system with a PepMap Neo C18 trap, 75um x 2cm (Thermo Scientific #164946) and EasySpray PepMap RSLC, 75um x25cm column (Thermo Scientific #ES902). LC-MS/MS was performed using Data Dependent Analysis on an Orbitrap Fusion MS system. A 180-minute gradient was applied at 250 nL/min, starting with 1% of solution B (90% ACN, 0.1% FA), and increasing to the final 35% of solution B. MS1 spectra were acquired at 120,000 resolutions with a scan range of 350-1500 m/z and MS2 were acquired in the ion trap with collision energy of 30. Raw data files were analyzed using Proteome Discoverer 2.4 (PD) with Sequest HT, searching the mouse protein database (UP000000589, downloaded March 30, 2021). Peptides identified with at least medium confidence were used for final results. Search parameters included two missed cleavages, a fragment mass tolerance of 0.5 Da, and a precursor mass tolerance of 10 ppm. Carbamidomethylation of cysteine was set as a fixed modification, while dynamic modifications included N-terminal acetylation, methionine oxidation, and SILAC labeling of Proline ^13^C_5_ for collagen synthesis or lysine ^13^C_6_^15^N_2_ and arginine ^13^C_6_ for global protein synthesis. Protein identifications were accepted at a false discovery rate (FDR) of 0.01.

### 2.7. Tracing of BCAA-Derived Branched-Chain Fatty Acids

To assess the incorporation of BCAAs into branched-chain fatty acids (BCFAs), WT and TAZ-KO cells were incubated with [U-^13^C_6_]-leucine, [U-^13^C_6_]-isoleucine, and [U-^13^C_5_]-valine as described in Sections 2.5.1 and 2.5.2. Following a 4-hour labeling period, cells were harvested, and lipids were extracted for analysis. The incorporation of ^13^C into the BCFAs isomyristic acid, isopalmitic acid, and isostearic acid was quantified by LC-MS at the Lipidomics Core Facility, Wayne State University.

### 2.8. Statistical analysis

Protein abundance ratios were calculated by dividing (Heavy) H-labeled by (Light) L-labeled protein abundance for each sample. Proteins were retained for analysis only if they had at least two non-missing abundance values in each experimental group (WT and TAZ-KO) and for H- and L-labeled abundances. Missing abundances were imputed by random forest imputation, performed separately for H- and L-labeled channels. Ratios were normalized to the sample median and log_2_-transformed. To assess global variation and group separation, group-level comparisons between WT (n = 4) and TAZ-KO (n = 4) were conducted using two-sample t-tests on median-normalized log_2_ H/L abundance ratios for each protein. Proteins were classified as differentially abundant if the TAZ-KO versus WT comparison of log_2_ H/L ratios met two criteria: an unadjusted p-value < 0.05 and an absolute fold-change > 1.5. All statistical and bioinformatics analyses for gene ontology were carried out with assistance from the Biostatistics and Bioinformatics Core at the Karmanos Cancer Institute.

### 2.9. Data availability

The mass spectrometry proteomics data have been deposited to the ProteomeXchange Consortium via the PRIDE [1] partner repository with the dataset identifier PXD078912 and 10.6019/PXD078912.

## 3. Results

### 3.1. TAZ-KO cells exhibit altered levels of BCAAs and proline and impaired glucose flux into NEAAs

PDH, the gatekeeper enzyme that channels pyruvate into the TCA cycle—is impaired in TAZ-KO cells as well as in TAZ-KO mouse cardiac tissue, marking PDH downregulation as a consistent feature of BTHS [7–9]. In parallel, TAZ-KO fibroblasts, TAZ-KO mouse heart mitochondria, and patient-derived iPSC cardiomyocytes show reduced expression of FAO genes and proteins [7, 11]. To explore whether AA metabolism is altered in TAZ-KO cells, we profiled steady state levels of intracellular AAs in WT and TAZ-KO cells using LC–MS. Levels of essential AAs such as BCAAs, lysine and threonine were reduced in TAZ-KO cells, while NEAAs showed mixed changes, with alanine and proline increased (Fig. 1 A and B).

**Figure 1:**
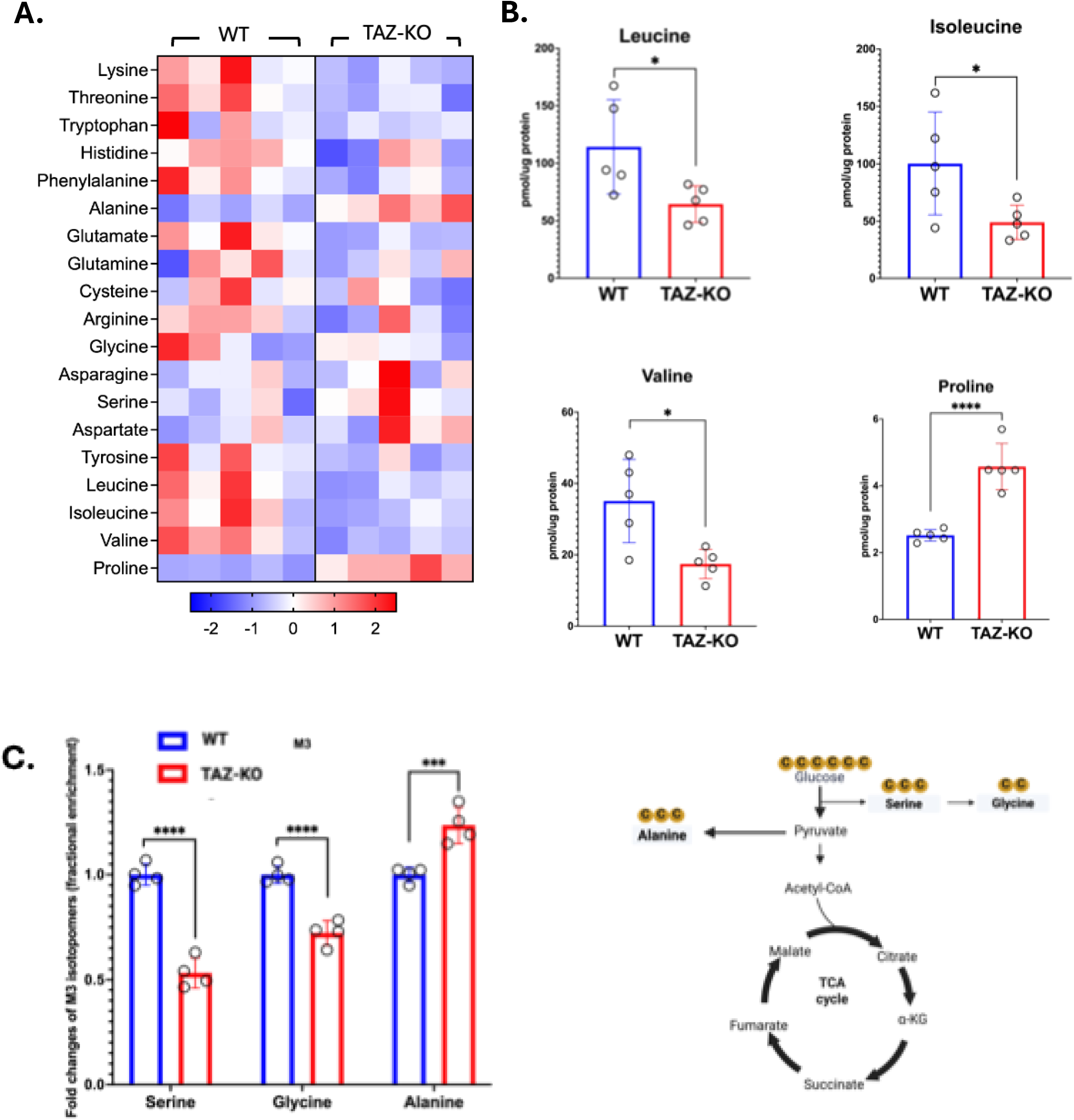
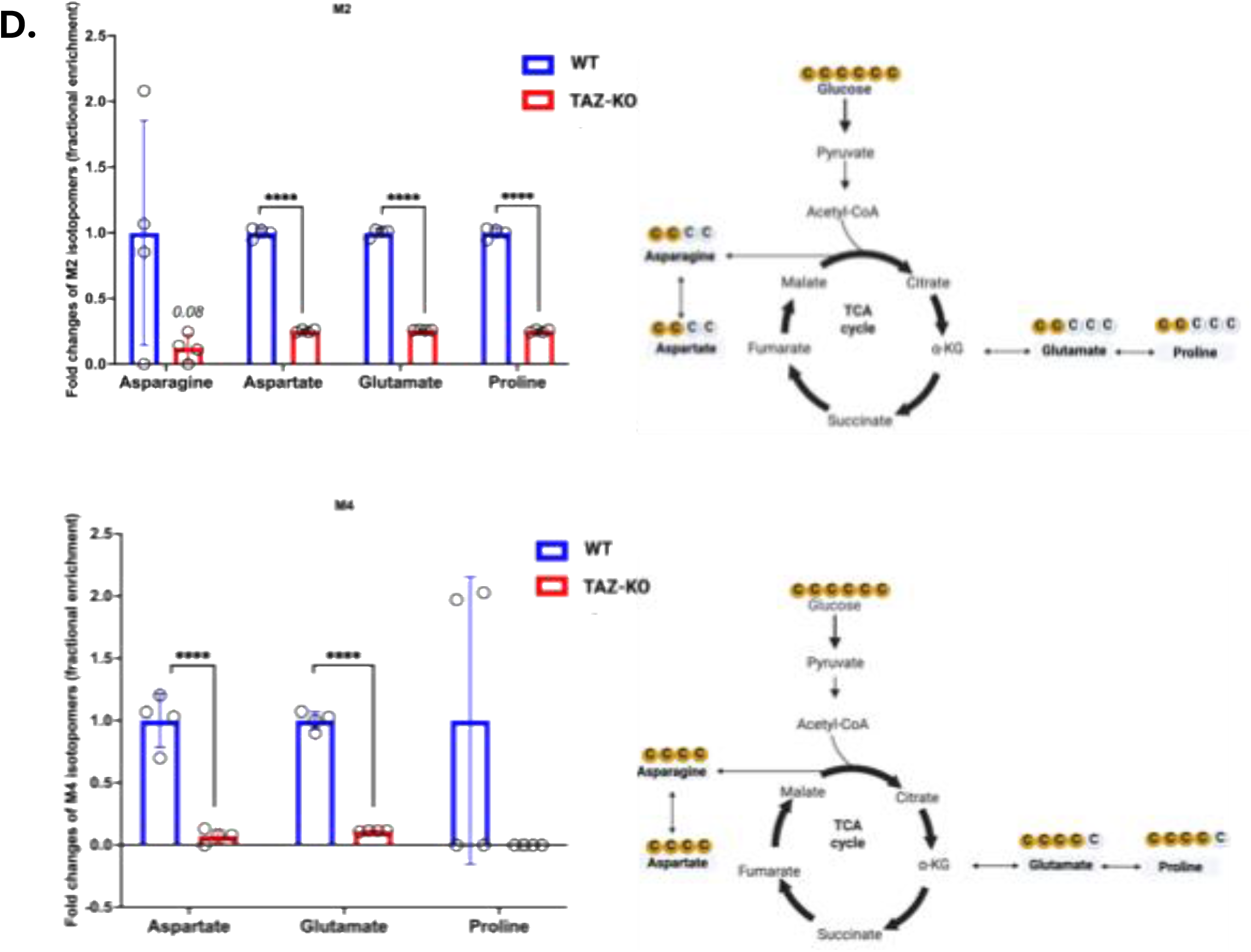
Altered amino acid metabolism and impaired glucose flux into NEAAs in TAZ-KO C2C12 myoblasts. **(A)** Amino acid levels in WT and TAZ-KO C2C12 myoblasts (N=5) as determined by LC-MS. Data are normalized to protein content and presented as a heat map auto-scaled to z scores and coded blue (low values) to red (high values). **(B)** The intracellular levels (pmol) of (BCAAs) leucine, isoleucine, valine, and proline were measured by LC-MS. Y-axis indicates steady-state intracellular AA levels, normalized to total protein (pmol/µg protein). Data shown are mean ± S.D. (N = 5). *, p < 0.05; ***, p < 0.001. **(C)** Mass isotopomers distribution (MID) of M3 isotopomers represents amino acids derived from glycolytic intermediates prior to conversion to two-carbon acetyl-CoA, and **(D)** MID of M2 (middle) and M4 (bottom) isotopomers represents amino acids derived from TCA cycle intermediates, as determined by LC-MS after 4-h incubation with [U-^13^C_5_]glucose. The X-axis represents amino acids derived from glycolytic and/or TCA cycle intermediates. The Y-axis represents fold change of fractional enrichment of isotopomers relative to WT. Data shown are mean ± S.D. (n = 4). **p < 0.001; ***p < 0.0001; ****p < 0.00001.

Since skeletal myopathy is one of the main clinical features of BTHS, we first aimed to gain insight into AA metabolism relevant to muscle composition and differentiation focusing first on BCAAs and proline. BCAAs comprise ∼35% of essential AAs in muscle proteins and are primarily catabolized in skeletal muscle (in contrast to AAs that are metabolized in the liver) [12–14]. Proline is critical for collagen synthesis and muscle differentiation [3, 19–22]. Levels of BCAAs (leucine, isoleucine, and valine) were decreased in TAZ-KO cells, with a 1.77-fold reduction in leucine and a 2-fold reduction in both isoleucine and valine. The reduction in intracellular BCAA levels could reflect increased catabolism, altered uptake, or changes in protein turnover. To investigate whether reduced BCAA uptake or altered catabolism contributes to these phenotypes, we performed stable isotope tracing and assessed transporter expression, as described later. We also observed 1.8-fold increased levels of proline, suggesting that entry of proline into the TCA cycle may be decreased (Fig. 1 B). Figure 1A also shows altered levels of other essential AAs (e.g. lysine, threonine, alanine, glutamate) in TAZ-KO cells. Glucose is the primary carbon source for most NEAAs (e.g. serine, alanine, glutamate, proline, asparagine, and aspartate), and our lab has previously demonstrated that glucose flux into the TCA cycle is impaired in TAZ-KO cells [7]. Among these, serine, threonine, and alanine are synthesized directly from glycolytic intermediates. [U-^13^C]glucose incorporation into most NEAAs was reduced in TAZ-KO cells relative to WT.

Serine, which is synthesized from 3-phosphoglycerate (3-PG), showed decreased M3-[^13^C]serine isotopomer enrichment in TAZ-KO cells, reflecting reduced glucose-derived carbon flux into serine biosynthesis (Fig. 1C). Glycine, which is derived from serine, showed decreased M2-[^13^C]threonine isotopomer enrichment from [U-^13^C]glucose. In contrast, alanine-synthesized from pyruvate via transamination-showed increased M3-[^13^C]alanine isotopomers enrichment in TAZ-KO cells (Fig. 1C). This is consistent with previously reported PDH downregulation in TAZ-KO cells, suggesting that impaired pyruvate entry into the TCA cycle may contribute to increased alanine synthesis via transamination, though further studies will be needed to establish a direct link.

Glutamate, proline, asparagine, and aspartate are derived from TCA cycle intermediates. M2 isotopomers are generated from intermediates in the first round of the TCA cycle, while M4 isotopomers arise from the condensation of M2-[^13^C]acetyl-CoA with M2-[^13^C]oxaloacetate in the second round. Both M2 (Fig. 1D, top panel) and M4 (Fig. 1D, bottom panel) isotopomers of glutamate, proline, asparagine, and aspartate were decreased in TAZ-KO cells relative to WT. These data demonstrate that glucose incorporation into NEAAs is impaired in TAZ-KO cells.

Collectively, these findings demonstrate that loss of TAFAZZIN broadly remodels amino acid metabolism, impairing glucose-derived carbon flux into NEAAs and potentially enhancing BCAA catabolism to replenish TCA cycle intermediates. Additionally, loss of TAFAZZIN increases proline catabolism via the TCA cycle.

### 3.2. BCAA transporter expression or uptake is not impaired in TAZ-KO cells

To determine whether reduced BCAA import contributes to the decreased intracellular BCAA levels observed in TAZ-KO cells, we assessed transporter expression and performed stable isotope tracing using [U-^13^C]leucine, [U-^13^C]isoleucine, and [U-^13^C]valine. Intracellular labeling efficiency and extracellular pool size of all three [^13^C]BCAAs were measured by LC-MS. Protein levels of LAT1 (SLC7A5), the primary plasma membrane BCAA transporter in skeletal muscle, and SLC25A44, the mitochondrial BCAA carrier, were unchanged in TAZ-KO cells (Supporting fig. 1A). Consistent with this, stable isotope tracing using [U-^13^C]-labeled BCAAs revealed effective intracellular labeling of all three BCAAs in both WT and TAZ-KO cells (Supporting fig. 1B). The fold change in extracellular [U-^13^C]leucine, [U-^13^C]isoleucine, and [U-^13^C]valine pool size was not significantly different between WT and TAZ-KO cells (Supporting fig. 1C), confirming that BCAA uptake was not impaired. These findings indicate that the reduced BCAA levels in TAZ-KO could be due to altered intracellular BCAA catabolism rather than impaired import.

### 3.3. BCAA catabolic enzymes are upregulated in TAZ-KO

BCAA catabolism is mediated by two key enzymes: BCAT (branched chain amino transferase), controlled at the expression level, and BCKDH (branched chain keto dehydrogenase), regulated by phosphorylation (Fig. 2A). During starvation, BCKDH activity increases markedly in muscle and heart, where AAs replace fatty acids (FAs) and ketone bodies as the main fuel, highlighting the role of BCKDH enzymatic activity in regulating BCAA catabolism [26]. BCKDH is inhibited by phosphorylation and reactivated by dephosphorylation [27–30] (Fig. 2A). To test if BCAA catabolism was increased in TAZ-KO, we measured mRNA and protein levels of BCAT2 and BCKDHE1a (phosphorylated and unphosphorylated) in WT and TAZ-KO myoblasts. While TAZ-KO cells showed slightly elevated mRNA expression of BCAT2 (Fig. 2B), BCAT2 protein levels were unchanged (Fig. 2C). In contrast, levels of BCKDHE1α mRNA and protein were elevated. Hyperphosphorylation of the regulatory E1α subunit of BCKDHE1α suppresses BCKDHE1α activity [28, 31–33]. Levels of the phosphorylated (inactive) protein were significantly elevated, suggesting that despite increased expression (Fig. 2C), phosphorylation may inactivate BCKDHE1α, creating a bottleneck that limits effective BCAA entry into the TCA cycle.

**Figure 2:**
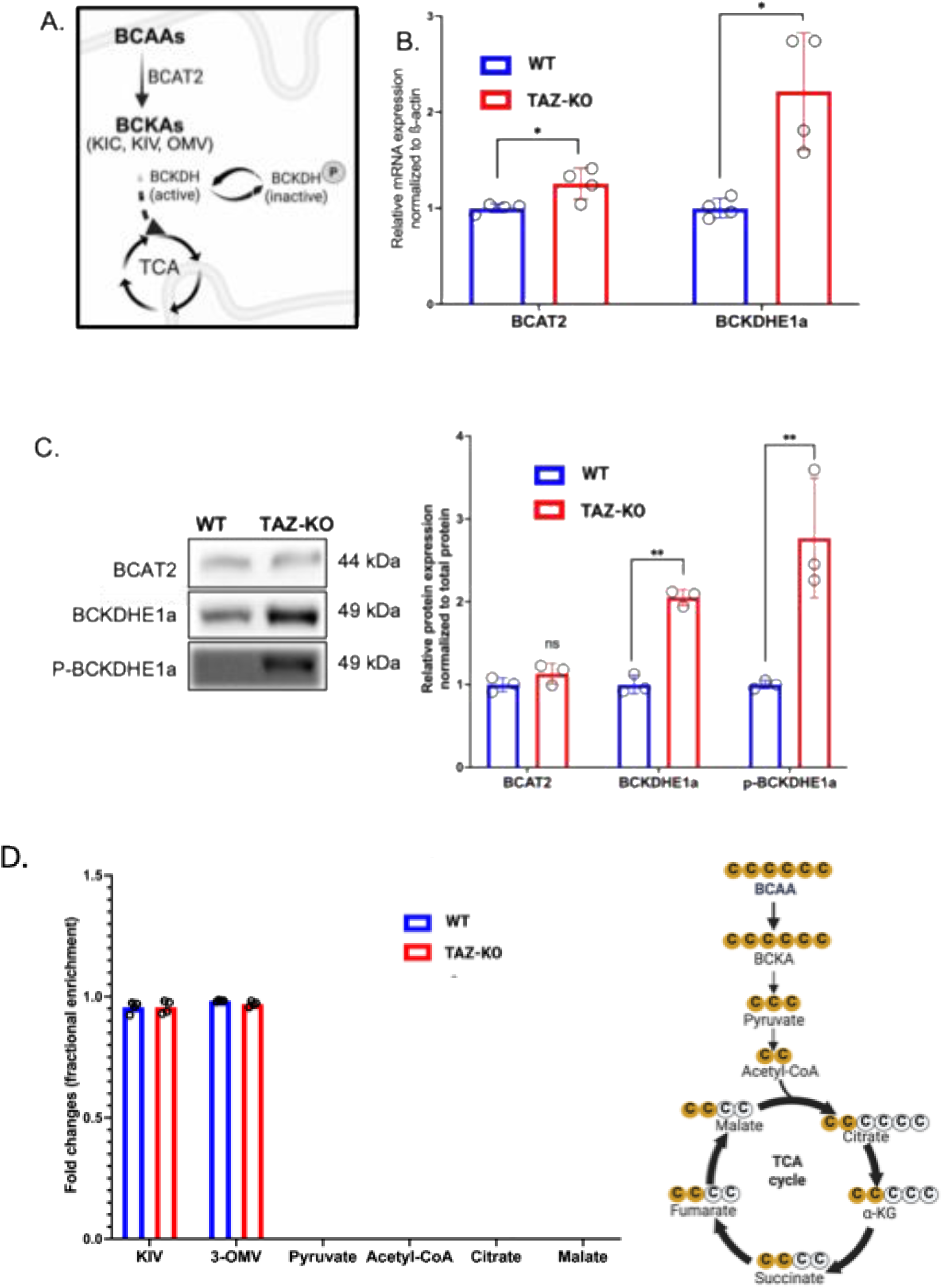
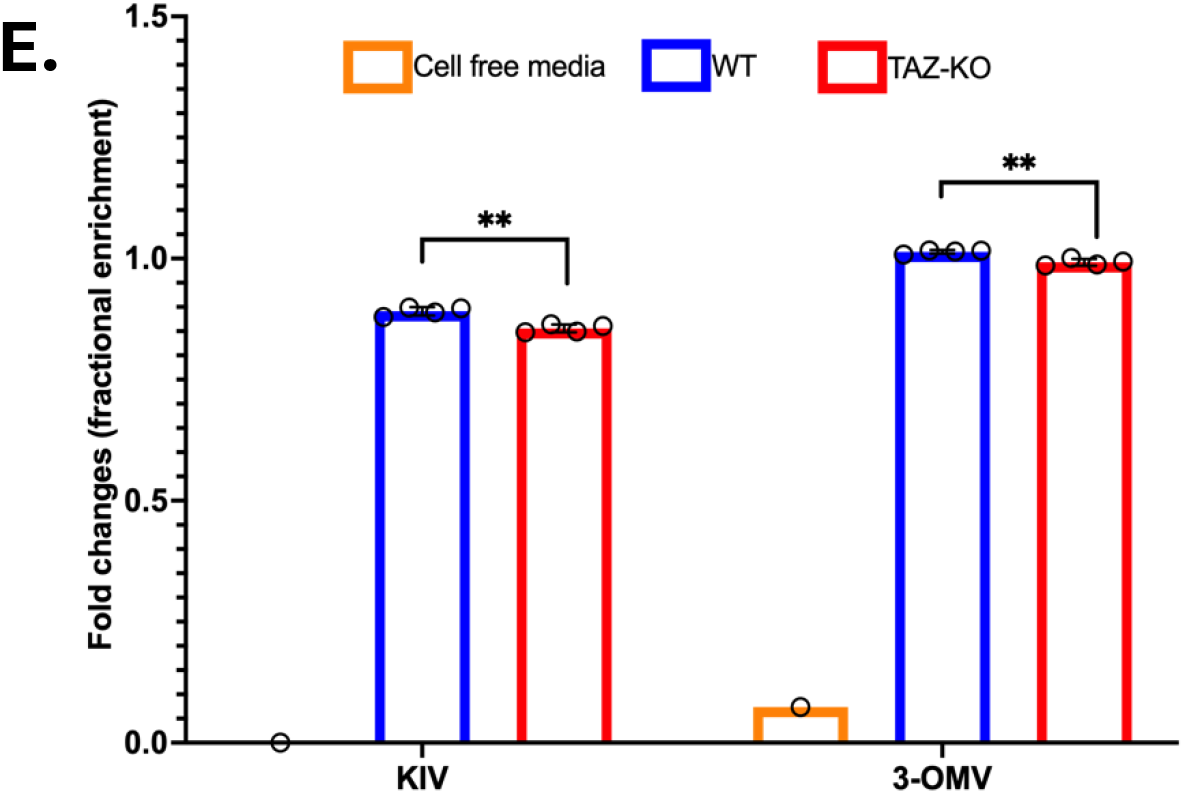
Loss of TAFAZZIN increases inhibitory BCKDH phosphorylation and impairs BCKA efflux. **(A)** Schematic overview of the catabolic pathway of BCAAs into the TCA cycle. Solid arrows denote single enzymatic reactions; dashed arrows indicate multi-step processes. **(B)** Transcript levels of key BCAA catabolic enzymes in WT and TAZ-KO myoblasts, assessed by qPCR and normalized to actin. **(C)** Protein expression of BCAT2, BCKDHE1α, and phospho-BCKDHE1α in WT and TAZ-KO myoblasts, assessed by western blot (n = 3). Left panel shows representative blots; right panel shows densitometric quantification normalized to total protein, performed using iBright analysis software. Statistical comparisons were made using unpaired, two-tailed Student’s t-tests. Data are presented as mean ±S.D. *p< 0.05; **p < 0.01. Primary antibodies: anti-BCAT2 (1:1000, 16417-1-AP, Proteintech), anti-BCKDHE1α (1:1000, 90198, Cell Signaling Technology), anti-phospho-BCKDHE1α (1:1000, 40368, Cell Signaling Technology). **(D)** Incorporation of [U-^13^C]BCAA-derived carbons into BCKAs and TCA cycle intermediates in WT and TAZ-KO C2C12 myoblasts, following 4-h incubation with [U-^13^C]leucine, [U-^13^C]isoleucine, and [U-^13^C]valine, as determined by LC-MS. X-axis represents BCKAs and TCA cycle intermediates; Y-axis represents fold change of fractional enrichment relative to WT. Data shown are mean ± S.D. (n = 4). **(E)** BCKAs detected in the culture media, indicating cellular export of BCKAs generated from labeled BCAA, from the same experiment described in (D). X-axis represents individual BCKAs (KIC, KIV, KMV); Y-axis represents fold change of fractional enrichment relative to WT. Cell-free blank media showed negligible enrichment, confirming BCKAs originated from cellular metabolism. Data shown are mean ± S.D. (n = 4).

To trace the metabolic fate of BCAAs into the TCA cycle in TAZ-KO cells, we used stable isotope tracing in collaboration with the Van Andel Institute Metabolomics Core. WT and TAZ-KO cells were cultured in high glucose (25 mM) DMEM supplemented with [U-^13^C]-labeled BCAAs- [U-^13^C_6_]leucine, [U-^13^C_6_]isoleucine, [U-^13^C_6_]valine- to monitor BCAA catabolism. BCAAs are first transaminated to BCAA-derived keto acids (BCKAs) such as-KIV (α-ketoisovalerate), and 3-OMV (3-oxomethylvalerate) (Fig. 2A). These BCKAs are then further processed by downstream enzymes into intermediates that can enter the TCA cycle. We observed no differences between WT and TAZ-KO cells with regard to incorporation of BCAAs into BCKAs-KIV (keto acid derived from valine) and 3-OMV (keto acid derived from isoleucine) (Fig. 3D). However, we observed no incorporation of labeled carbon into downstream TCA cycle intermediates (Fig. 2D). To investigate the fate of BCKAs, given the absence of incorporation into the TCA cycle, we examined the culture media from WT, TAZ-KO, and cell-free controls. [U-^13^C_5_]BCKAs were detected in the media of both WT and TAZ-KO cells, while cell-free media contained only negligible levels, confirming that BCKAs originated from cellular metabolism (Fig. 2E). Notably, TAZ-KO cells exported fewer BCKAs than WT, suggesting that BCKAs are rerouted toward an alternative fate rather than being exported out or diverted to the TCA cycle.

**Figure 3:**
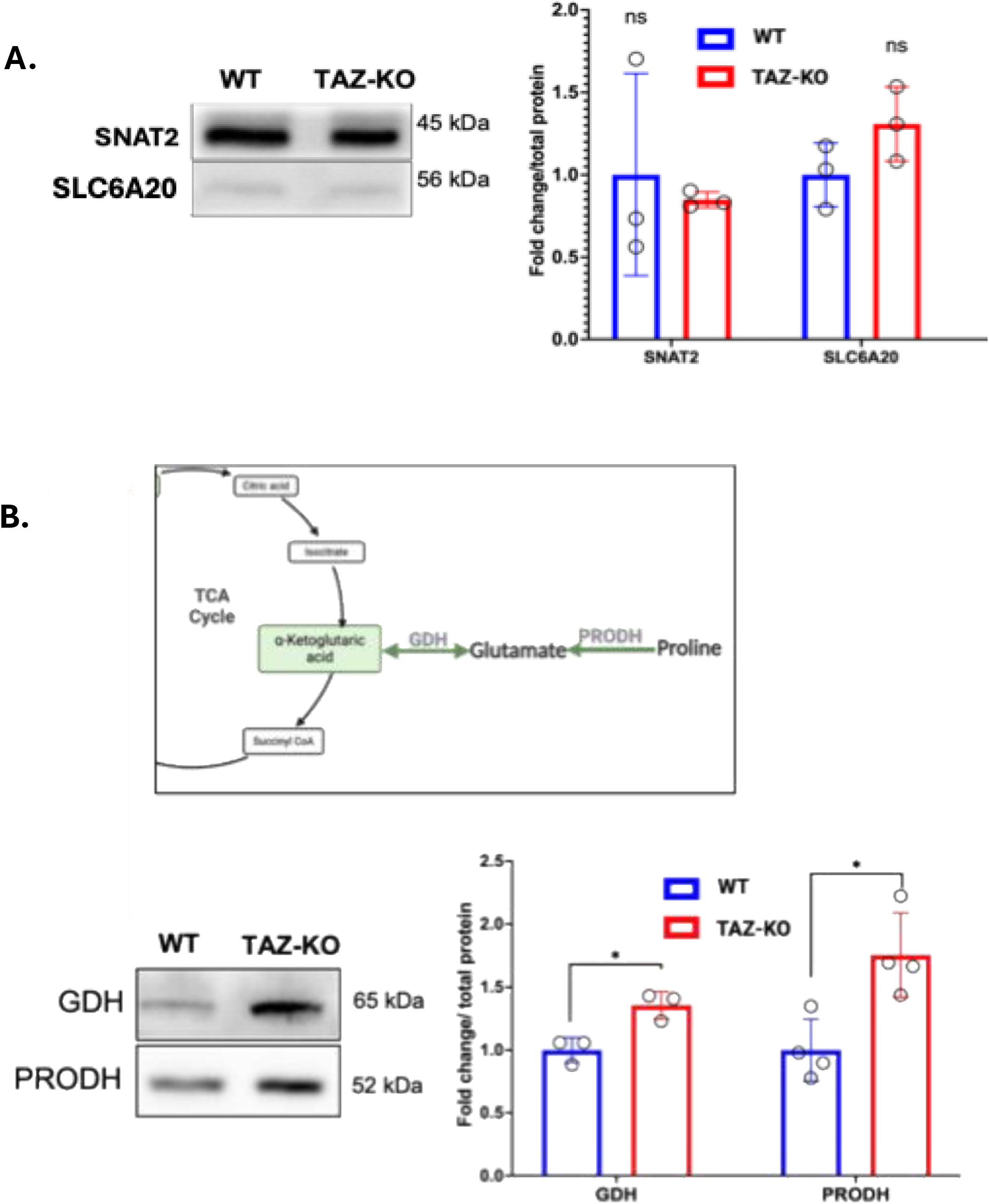
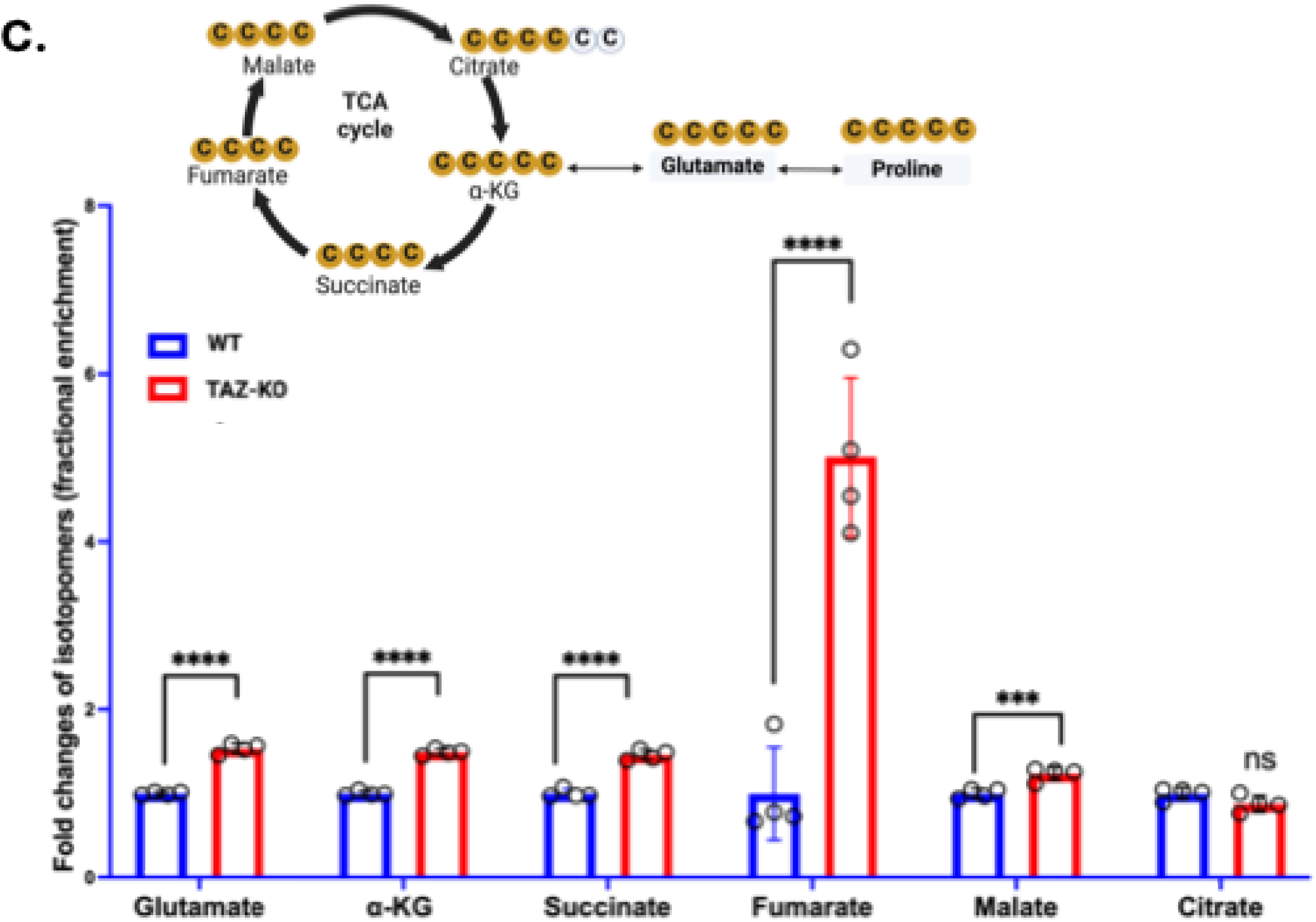
Increased expression of proline catabolic enzyme and increased incorporation of [U-^13^C_5_]-proline into TCA cycle intermediates in TAZ-KO cells. **(A)** Protein levels of proline transporters were assessed by western blot analysis of WT and TAZ-KO cell lysates. Left panel shows representative blot images. Densitometric analysis was performed using iBRIGHT (Thermo Fisher) software, and protein levels were normalized to total protein (right panel). Y-axis represents fold changes in relative protein expression. Data are presented as mean ± S.D. (N=3). Statistical comparisons were made using unpaired, two-tailed Student’s t-tests. Primary antibodies: anti-SNAT2 (1:1000, 25928-1-AP; Proteintech) and anti-SLC6A20 (1:1000, 30843-1-AP; Proteintech). **(B)** Schematic depicting proline catabolism (top panel): proline is converted to glutamate by PRODH, which is subsequently converted to alpha-ketoglutarate by GDH. Protein expression of PRODH and GDH was assessed by western blotting of WT and TAZ-KO myoblast lysates. Data are presented as mean ± S.D. (n=3). *p < 0.05. Primary antibodies: anti-GDH (1:1000, D9F7P; Cell Signaling) and anti-PRODH (1:2000, 22980-1-AP; Proteintech). **(C)** WT and TAZ-KO cells were incubated with [U-^13^C₅]-proline for 8 hours, and intracellular metabolites were analyzed by LC-MS. MID of M5 isotopomers represents glutamate and alpha-ketoglutarate retaining all five labeled carbons from proline. MID of M4 isotopomers represents downstream TCA cycle intermediates. Y-axis shows fold changes in fractional enrichment relative to WT, normalized to cell number. Data are presented as mean ± S.D. (n=4). **p < 0.01, ***p < 0.001, ****p < 0.0001.

To investigate the fate of BCKAs that were not exported into the media, given the reduced BCKA efflux observed in TAZ-KO relative to WT, we traced the incorporation of [U-^13^C]-labelled BCAAs into branched-chain fatty acids (BCFAs) (e.g. isopalmitic acid and isostearic acid). No difference in [U-^13^C]BCAA incorporation into BCFAs was observed between WT and TAZ-KO (Supporting Information 2). Collectively, these results suggest that under high-glucose conditions, BCAA catabolism is truncated at the level of BCKA oxidation in both WT and TAZ-KO cells. In TAZ-KO cells, this truncation is accompanied by reduced BCKA efflux into the extracellular media, rather than increased entry into the TCA cycle for energy production. Furthermore, the reduced BCKA efflux was not explained by increased incorporation into BCFAs.

### 3.4. Levels of proline transporters are unaltered in TAZ-KO cells

To test whether the proline increase results from enhanced import or cytosolic synthesis, we quantified protein levels of the proline transporters SIT1 (Sodium/Imino-acid Transporter 1), encoded by SLC6A20A, and SNAT2 (Sodium-coupled neutral amino acid transporter 2), encoded by SLC38A2, which are primarily responsible for proline transport in the murine brain and mammary gland. Both transporters are regulated by their expression levels [34–37]. We observed no significant differences between WT and TAZ-KO cells in protein expression of SNAT2, or SLC6A20 (SIT1) (Fig. 3A). This suggests that the elevated proline levels in TAZ KO are not due to altered transporter or importer expression but instead arise from altered proline metabolism.

### 3.5. TAZ-KO cells exhibit increased flux of proline into the TCA cycle

Proline is a NEAA that can be synthesized from glutamine or TCA cycle intermediates. A previous report suggested that flux from glutamine to proline was increased in TAZ deficient mouse cardiac tissue [17]. However, elevated levels may also result from impaired utilization. To test if proline catabolism into the TCA cycle is altered, we examined levels of proline dehydrogenase (PRODH), which converts proline to glutamate in mitochondria, and glutamate dehydrogenase (GDH), which converts glutamate to α-ketoglutarate (α-KG) (Fig. 3B, top panel schema). Both enzymes exhibited increased protein levels in TAZ-KO cells (Fig. 3B, bottom panel), suggesting that proline metabolism into the TCA cycle may be increased in these cells.

To test whether proline metabolism into the TCA cycle is altered, we supplemented cells with [U-^13^C_5_]proline and traced its incorporation into TCA cycle intermediates using LC-MS. Surprisingly, M5-[^13^C]proline flux into M5-[^13^C]glutamate and all TCA cycle intermediates, including M5-[^13^C]*α*-KG, M4-[^13^C]succinate, M4-[^13^C]fumarate, M4-[^13^C]malate and M4-[^13^C]citrate were markedly elevated in TAZ-KO cells (Fig. 3C). As previously reported, glucose-derived carbon flux into the TCA cycle is reduced in TAZ-KO cells, the elevated proline flux into TCA cycle intermediates therefore likely reflects increased anaplerosis to compensate for this deficit.

### 3.6. Collagen synthesis is impaired in TAZ-KO cells

Increased incorporation of proline into TCA cycle intermediates (Fig. 5) suggests that proline may be preferentially directed toward catabolism rather than collagen synthesis in TAZ-KO cells [38]. To address this possibility, we examined collagen gene expression, collagen content, and proline hydroxylation in TAZ-KO cells. Collagen supports ECM structure, cell adhesion, and muscle differentiation; therefore, impaired collagen synthesis or organization may impair myogenesis and contribute to the skeletal myopathy seen in BTHS [18–22, 25].

Type I collagen is the most prevalent form of collagen protein, accounting for about 90% of total collagen. It provides structural strength and support to connective tissues [39]. Type VI collagen plays diverse roles in tissues, including providing structural support in the ECM, protecting against apoptosis and oxidative stress, and regulating cell differentiation [40]. RNA-seq analysis comparing TAZ-KO and WT C2C12 undifferentiated myoblasts reported previously [41] revealed that genes encoding Type I and Type VI collagen were significantly downregulated in TAZ-KO cells (Fig. 4A). In agreement with this, expression of the Type I collagen protein Col1a1 was undetectable in TAZ-KO, suggesting impairment of collagen production (Fig. 4B). This indicates that TAZ deficiency disrupts collagen gene expression, particularly Type I and Type VI, which are critical for structural support and cell adhesion. The downregulation of Col1a1 expression further suggests that defective ECM remodeling may be a key mechanism contributing to impaired muscle differentiation and skeletal myopathy in BTHS. Consistent with decreased expression of these genes, we observed a significant reduction in total collagen content in TAZ-KO cells (Fig. 4C) as reflected in markedly decreased levels of hydroxyproline, a collagen-specific AA comprising ∼13% of total collagen weight [21, 22]. Analysis of the [U-^13^C₅]-proline tracing data revealed a significant decrease in ^13^C5-labeled hydroxyproline pool size in TAZ-KO cells compared to WT (Fig. 4D), with unchanged P4H protein expression (Supplementary Fig. 4K). This suggests that in TAZ-KO cells, proline is preferentially catabolized te replenish TCA cycle intermediates rather than being channeled toward P4H-mediated hydroxylation, resulting in reduced hydroxyproline production.

**Figure 4:**
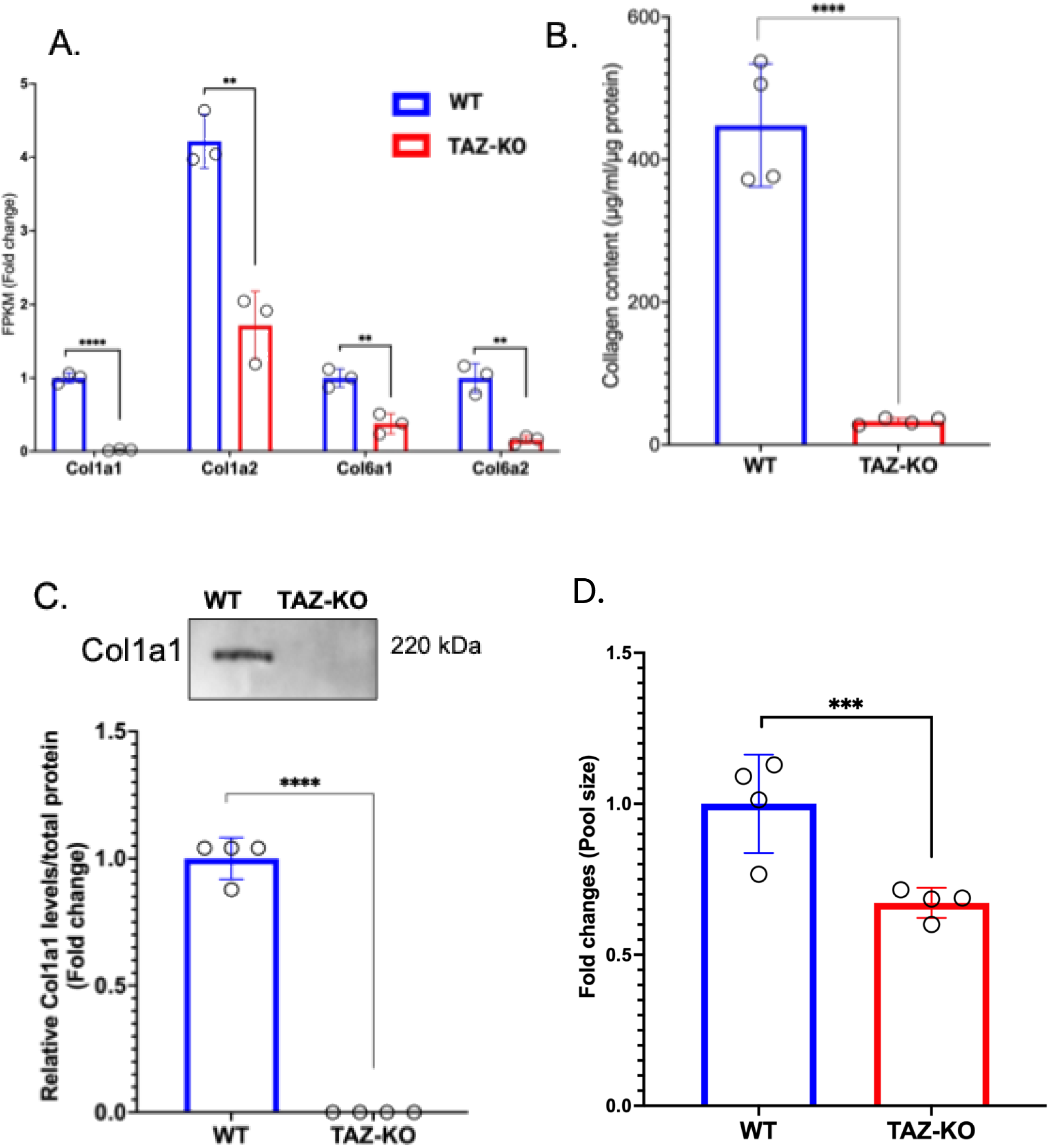
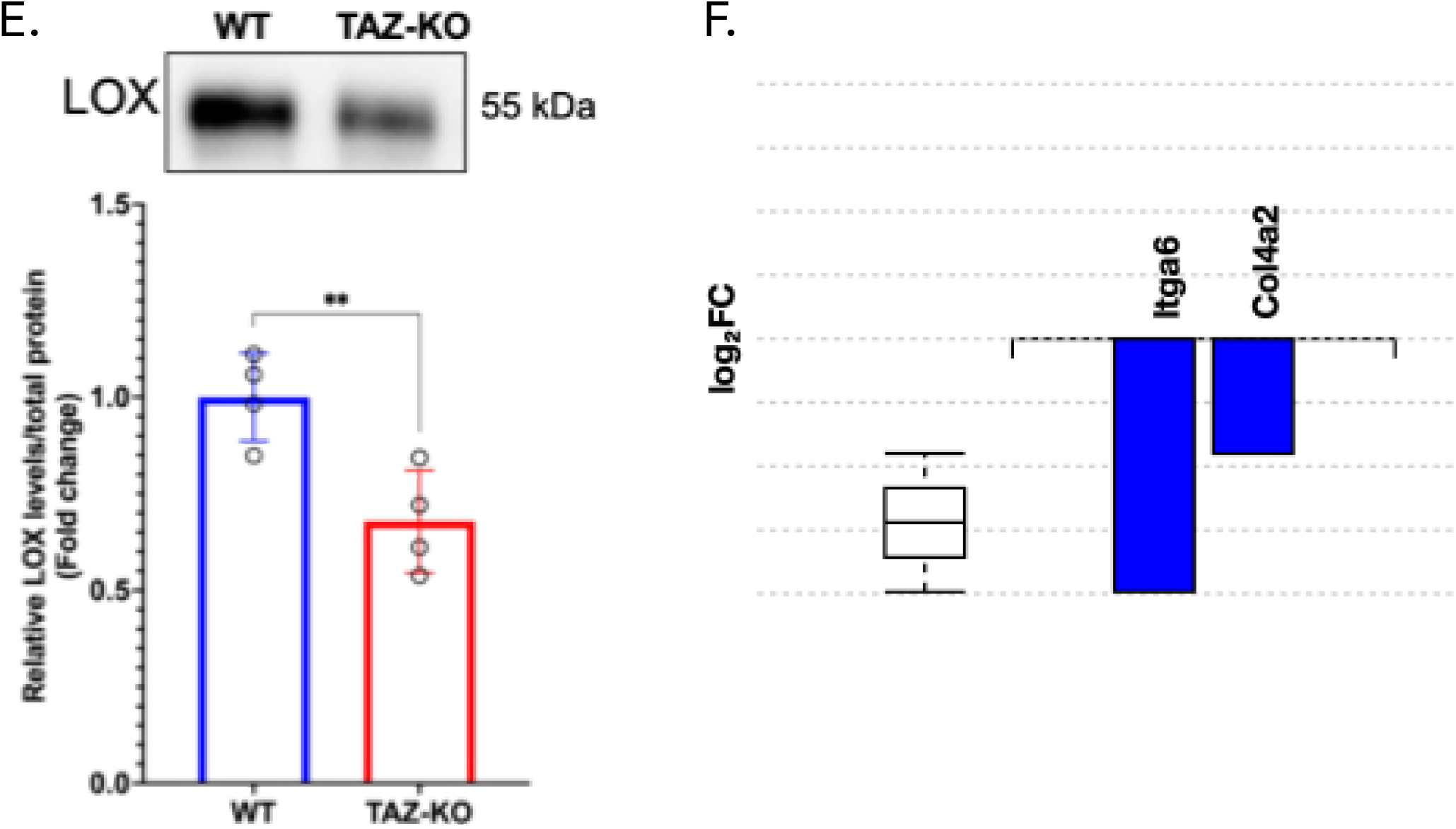
Reduced collagen content and collagen crosslinking enzyme expression in TAZ-KO cells. **(A)** RNA-seq analysis of Type I and VI collagens. Fragments per kilobase million (FPKM) values obtained from RNA-seq analysis were used to quantify the expression of collagen type I and VI isoforms in WT and TAZ-KO myoblasts. The Y-axis represents fold changes in FPKM values. **(B)** Collagen levels were measured in WT and TAZ-KO cell lysates by quantifying hydroxyproline content (μg of hydroxyproline/ml in Y-axis). Data are presented as mean ± SD (N = 4, with three technical replicates per sample. **(C)** Representatives immunoblot images (top) of Col1a1 in WT and TAZ-KO with corresponding quantification (bottom). **(D)** Fold change of total pool size of hydroxyproline in WT and TAZ-KO cells incubated with [U-^13^C₅]-proline for 8 hours, measured by LC-MS. Y-axis represents fold change in hydroxyproline pool size relative to WT. Data are presented as mean ± S.D. (n=4). **p < 0.001, unpaired two-tailed Student’s t-test. **(E)** Representatives immunoblot images of LOX protein levels in cell lysates (N = 3) (top). Band intensities were quantified by densitometry with ibright analysis software (bottom). Statistical analyses were performed using an unpaired, two-tailed Student’s t-test. Data are presented as mean ± S.D. (**p < 0.01, ***p < 0.001, ****p < 0.0001). Primary antibodies used were anti-Col1a1 (1:1000, 72026; Cell signaling technology) and anti-LOX (1:1000, 58135; Cell signaling technology). **(E)** Bar plot showing differential gene expression in TAZ-KO myoblasts relative to WT (N=4) measured by protein synthesis using [U-^13^C_5_,^15^N_2_] - Lys and [U-^13^C_6_] - Arg quantified by LC–MS. Genes with an absolute fold change >1.5 and an adjusted p-value <0.05 were considered significant. Gene ontology analysis of downregulated genes was performed using iPathwayGuide. Y-axis represents log_2_ fold change of differentially expressed genes.

The synthesis of collagen starts in the ER as pro-collagen, where it undergoes key post-translational modifications: removal of the N-terminal signal peptide, hydroxylation of lysine and proline, and glycosylation of specific hydroxylysines. Three modified procollagen chains twist into a triple helix, thereby forming tropocollagen, and then move to the Golgi apparatus for final processing and secretion. In the cellular space, lysyl oxidase (LOX) catalyzes covalent crosslinks between tropocollagen molecules, forming stable collagen fibrils. We assessed the expression of the LOX enzyme and found its protein levels were significantly decreased in TAZ-KO cells (Fig. 4E).

To measure the rate of collagen protein synthesis, WT and TAZ-KO cells initially cultured in “light” medium containing unlabeled Lys and Arg were switched to “heavy” medium supplemented with labeled [U-^13^C_6_, ^15^N_2_]-Lys and [U-^13^C_6_]-Arg for 24 hours. Incorporation of these heavy isotopes enabled discrimination between newly synthesized and pre-existing proteins. Using SILAC analysis with ^13^C-labeled Lys and Arg, proteomics data revealed that the rate of newly synthesized Type IV collagen was significantly reduced (Fig. 4F). Integrin alpha 6 (*Itga6*), a cell surface receptor which is essential in cell adhesion, signaling, and interaction with the extracellular matrix (ECM), was also significantly reduced in TAZ-KO cells (Fig. 4F). This suggests that the synthesis of new collagen and cell adhesion proteins and the processing of collagen in the ECM are impaired in TAZ-KO cells.

### 3.7. Impaired proline incorporation into collagen and ECM proteins in TAZ-KO

To evaluate the incorporation of proline into newly synthesized ECM proteins, we supplemented cells with [U^13^C₅]proline and quantified its incorporation into collagen and ECM remodeling proteins by LC-MS using SILAC. A total of 6,767 proteins were identified from eight samples (four WT and four TAZ-KO), and 1,380 proteins were retained for downstream analysis. Volcano plot analysis revealed 72 genes upregulated (red circles) and 126 genes downregulated (blue circles) in TAZ-KO cells (|fold change| >1.5, unadjusted p-value <0.05) (Fig. 5A). Gene ontology (GO) cellular component analysis (classifies genes based on their subcellular or extracellular localization) indicated that the most strongly affected category in TAZ-KO cells was the cell periphery, while GO biological pathway analysis (classifies genes based on their functional pathways and activities) showed marked suppression of ECM remodeling pathways (Fig. 5B). Figure 5C provides a schematic overview of key ECM and collagen-associated proteins. 11 ECM-related genes - Perlecan (HSPG2), PHLDB1, CLASP1, Flotillin-1 (FLOT1), HSP47 (SERPINH1), Fibronectin-1 (FN1), SPARC, Cathepsin L (CTSL), Glypican-1 (GPC1), HTRA1, and Tensin-2 (TNS2) - were significantly downregulated in TAZ-KO cells (highlighted as enlarged, solid circles and labeled with their respective symbols) (Fig. 5A). Gene ontology (GO) analysis further demonstrated that these downregulated ECM genes clustered into three major processes: collagen metabolism, collagen-containing ECM, and ECM organization (Fig. 5C). Taken together, these findings indicate that loss of TAZ impairs proline incorporation into newly synthesized collagen and ECM proteins, suggesting a broad suppression of ECM remodeling and organization.

**Figure 5:**
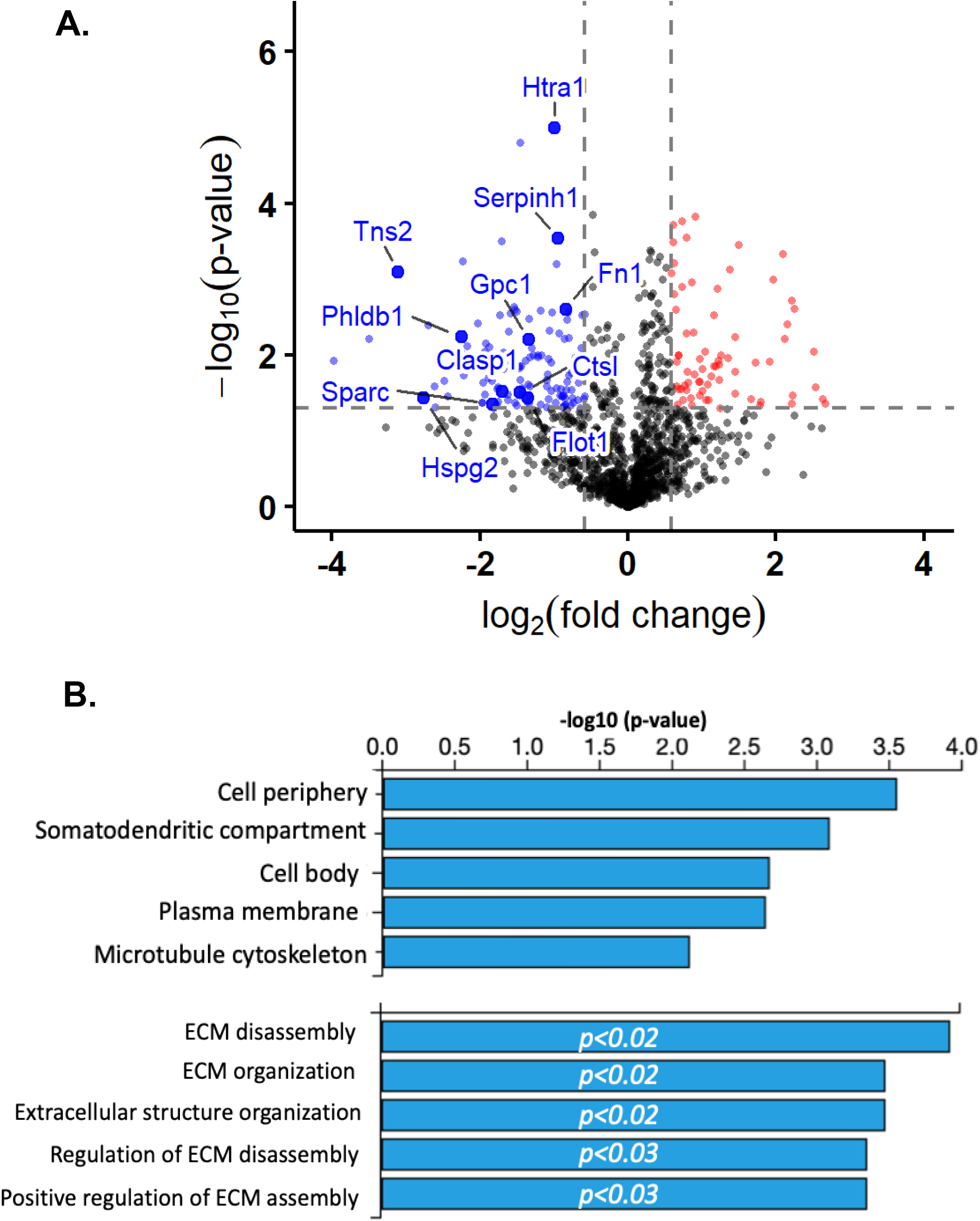
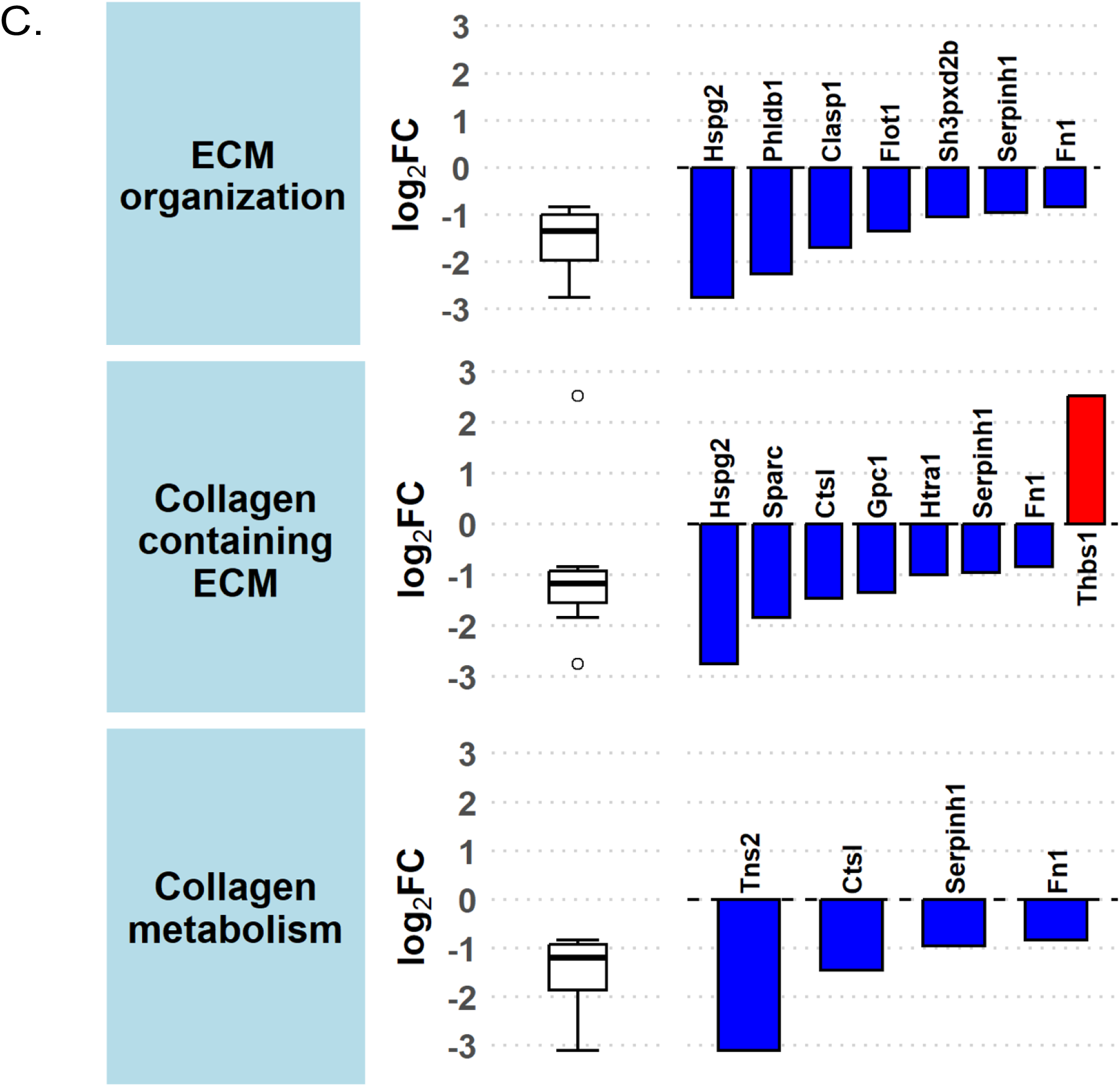
Decreased incorporation of [U-^13^C_5_]-proline into collagen and ECM proteins. (**A)** Volcano plot of the significant differentially expressed genes between the WT and TAZ-KO. Genes with an absolute fold change >1.5 and an unadjusted p-value <0.05 were classified as significant. X-axis represents the log_2_ fold change and Y-axis represents −log₁₀ p-value of differentially expressed genes. (**B**) GO analysis of significantly downregulated genes related to ECM remodeling and organization, performed with iPathwayGuide software. **(C)** GO analysis of downregulated genes revealed suppression of specific genes associated with ECM-related pathways, including collagen metabolism, collagen-containing matrix, and ECM organization. Y-axis represents the log_2_ fold change of differentially expressed genes.

## 4. Discussion

In this study, we demonstrate that loss of TAZ disrupts amino acid metabolism and collagen biosynthesis. TAZ-KO cells showed reduced levels of BCAAs, reduced glucose flux into amino acids, and increased levels of proline (Fig. 1), accompanied by increased expression of proline catabolic enzymes (Fig. 3B) and enhanced flux of proline into TCA cycle intermediates (Fig. 3C). Despite elevated intracellular proline, collagen synthesis was markedly reduced, as evidenced by decreased proline hydroxylation and collagen content (Fig. 4 B-C), decreased expression of collagen isoforms (Fig. 4A) and crosslinking enzymes (Fig. 4D), and diminished incorporation of proline into collagen and ECM proteins (Fig. 5). Our findings identify a previously unrecognized role for TAZ in coordinating AA metabolism with ECM remodeling.

Previous studies demonstrated that glucose and FA metabolism are altered in TAZ-KO cells [7, 9] [11], suggesting that TAZ-deficient cells may exhibit increased reliance on AAs for energy metabolism. In the current study, we found that AA metabolism is reprogrammed, as evidenced by a significant reduction of BCAAs, lysine and threonine, and elevated levels of alanine and proline in TAZ-KO cells (Fig.1). This metabolic shift suggests a compensatory mechanism aimed at supporting energy production in TAZ-deficient cells, with BCAAs and proline playing a critical role in muscle metabolism in BTHS.

Overexpressing BCAT-encoding genes *BAT1* or *BAT2* in *taz1*Δ yeast cells has been shown to restore oxygen consumption to WT levels, emphasizing the importance of BCAT-mediated BCAA metabolism in maintaining mitochondrial bioenergetics [42]. Elevated levels of BCKDH phosphorylation were observed in TAZ-KO cells, suggesting a bottleneck in downstream oxidation, preventing full utilization of BCAAs for energy production. In mice, high glucose suppresses the expression of enzymes involved in the BCAA degradation pathway [43]. Under high-glucose conditions, BCAA incorporation into TCA cycle intermediates was not detected in either WT or TAZ-KO cells, as BCAA catabolism appeared to halt at the BCKA formation step (Fig. 2D). Interestingly, fractional enrichment of [U-^13^C]-BCKAs in the WT and TAZ-KO media was higher than in cell-free media controls, indicating that C2C12 cells actively export BCKAs. However, TAZ-KO cells exhibited lower BCKA enrichment in the media relative to WT, suggesting impaired BCKA export. Studies have shown that endogenous branched chain fatty acids (BCFA) can be synthesized from BCAA precursors through the BCAA catabolic pathway [44]. However, BCAA isotope tracing did not reveal any detectable difference in intracellular BCFA labeling between WT and TAZ-KO cells (Supporting info. 2), indicating that reduced BCKA efflux in TAZ-KO cells is not explained by increased shuttling into BCFA synthesis. The metabolic fate of the missing BCKA pool therefore remains unresolved. However, we cannot rule out subtle differences in BCAA export or increased incorporation into newly synthesized proteins as contributing factors to the reduced intracellular BCAA levels observed in TAZ-KO cells. Together, these findings highlight the complexity of BCAA metabolism in TAZ-KO and underscore the need for further investigation to fully establish the mechanistic basis of these observations.

In contrast to the reduced BCAA levels observed in TAZ-KO cells, proline was significantly elevated (Fig. 1A&B). To investigate whether this accumulation reflected altered proline catabolism, we examined the expression of proline catabolic enzymes and traced the flux of [U-^13^C_5_]proline-derived carbon into TCA cycle intermediates. Proline, a NEAA derived from glutamine and glutamate, replenishes TCA cycle intermediates via two key enzymes, GDH and PRODH. Previous studies have shown that when glucose is depleted or its metabolism impaired, cells become entirely reliant on GDH for growth [45], and that PRODH is upregulated to help tumor cells survive under metabolic stress [46].

In TAZ-KO cells with impaired glucose and FA metabolism, we observed elevated expression of both enzymes that enable proline to enter the TCA cycle. Additionally, we detected increased incorporation of [U-^13^C_5_]proline into glutamate, α-ketoglutarate, and other TCA cycle intermediates. Our findings suggest that proline compensates for TCA cycle disruption in TAZ-KO cells through anaplerotic pathways.

Beyond BCAAs and proline, steady state levels of other NEAAs were also altered in TAZ-KO cells (Fig. 1A). Myoblasts rely on glucose for energy production. Glucose also feeds NEAA biosynthesis through glycolytic and TCA cycle intermediates. Given the reduced glucose flux into the TCA cycle in TAZ-KO cells [7, 47], we asked whether glucose flux to NEAA biosynthesis was affected. [U-^13^C]glucose tracing revealed, glucose flux to serine and glycine (synthesized from the glycolytic intermediate 3-phosphoglycerate) was markedly reduced in TAZ-KO cells (Fig. 1C), whereas their total intracellular pools were unchanged (Supporting info. 3A). The source responsible for maintaining the pools was not determined in this study; however, uptake of exogenous serine and glycine is likely to be the contributor [48, 49]. On the other hand, glucose flux to alanine was increased (Fig. 1C), consistent with reduced pyruvate entry into the TCA cycle through reduced PDH activity in TAZ-KO cells [7] and the rerouting of pyruvate toward alanine synthesis. Despite this increased glucose flux, the total alanine pool was reduced (Supporting info. 3A), which suggests an increased conversion of pyruvate to alanine accompanied by increased alanine utilization and export [50, 51], although these also require further validation.

NEAAs such as aspartate, glutamate, proline, and asparagine are derived from TCA cycle intermediates. We observed that glucose flux into aspartate, glutamate and proline was reduced, whereas asparagine labeling was unchanged (Fig. 1D). Total pool of aspartate and glutamate were unchanged, while proline and asparagine were increased (Supporting info. 3B). Together, these findings indicate reduced glucose flux into TCA cycle-derived NEAAs in TAZ-KO cells, with their pools maintained by an alternative carbon source, most likely glutamine. In TAZ deficient mouse hearts and patient-derived cardiomyocytes, glutamate uptake is increased and replenishes the TCA cycle, while elevated expression of asparagine synthetase (ASNS) and the proline synthetic enzymes P5CS (ALDH18A1) and PYCR1 upregulate asparagine and proline synthesis [17]. Our study demonstrated also that proline is catabolized to replenish TCA cycle intermediates in TAZ-KO cells (Fig. 3C) and may also indicate altered synthesis of proline-rich proteins (e.g. collagen) [3, 19–22]. Together, these findings indicate that loss of TAFAZZIN reroutes NEAA synthesis away from glucose, highlighting disrupted coordination between energy and AA metabolism.

Proline is a key component of collagen and the primary structural protein of the ECM [21, 22]. The increased incorporation of proline into the TCA cycle for replenishment (Fig. 3C) suggested that collagen biosynthesis is affected in TAZ-KO cells. The extracellular proline level significantly impacts collagen biosynthesis, highlighting the importance of proline availability for collagen synthesis rates [21]. Consistent with this, TAZ-KO cells exhibited reduced total and intracellular collagen levels, along with reduced collagen crosslinking enzyme LOX, indicative of impaired collagen synthesis (Fig. 4C-D). In agreement with this, the SILAC tracing exhibited reduced incorporation of proline into newly synthesized proteins associated with collagen metabolism, collagen-containing ECM, and ECM organization (Fig. 5). Collagen, the most abundant ECM protein, is essential for tissue integrity and force transmission during myotube formation [19, 20]. Because collagen is highly enriched in proline, disruptions in proline metabolism or collagen synthesis can impair ECM remodeling, leading to defective differentiation and muscle fiber maturation [18, 21, 22]. Our findings suggest that impaired collagen synthesis and reduced proline flux into collagen and other ECM proteins may link TAZ loss to skeletal myopathy in BTHS.

Collectively, this study reveals a novel mechanistic link between mitochondrial lipid remodeling, metabolic regulation, and ECM biology, suggesting that loss of TAZ drives metabolic reprogramming that could contribute to muscle pathology.

## Supporting information

Supplemental informations

## Acknowledgements

This work was supported by National Institutes of Health grants HL117880 and HL174611 (to M.L.G.). Stable isotope tracing metabolomics experiments were conducted by the Van Andel Institute Mass Spectrometry Core (RRID:SCR_024903). We also acknowledge the assistance of the Wayne State University Proteomics Core that is supported through NIH grants P30ES036084, P30CA022453 and S10OD030484. The Biostatistics and Bioinformatics Core is supported, in part, by NIH Center grant P30 CA022453 to the Karmanos Cancer Institute at Wayne State University

