## Supplemental informations for "Loss of TAFAZZIN leads to perturbation of amino acid metabolism and reduction of collagen synthesis"

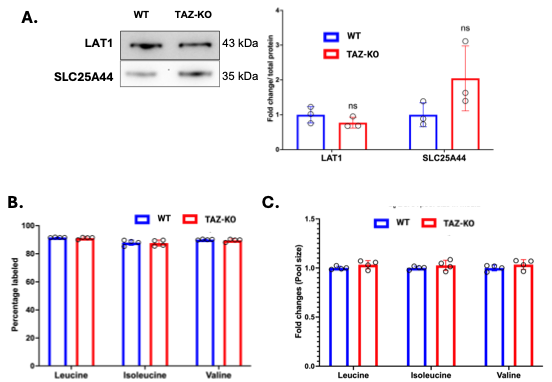

**Supporting info 1: TAZ deficiency does not impair BCAA transporter expression or uptake.** **(A)** Protein levels of BCAA transporters were assessed by western blot analysis of WT and TAZ-KO cell lysates. Left panel shows representative blot images. Densitometric analysis was performed using iBRIGHT (Thermo Fisher) analysis software. Protein levels were normalized to total protein (right panel). Y-axis of the bar graphs in the right panels represents fold changes in relative protein expression, normalized to total protein content. Statistical comparisons were made using unpaired, two-tailed Student's t-tests. Data are presented as mean ± S.D. (N=3). BCAA uptake was assessed by measuring intracellular labelling (B) and extracellular (C) BCAA pool size following 4 hours incubation of WT and TAZ-KO C2C12 myoblasts with [U-¹³C]-Leucine, [U-¹³C]-Isoleucine, and [U-¹³C]-Valine added together to the culture media, as determined by LC-MS. **(B)** Y- axis of the bar graph shows the percentage of intracellular leucine (M+6), isoleucine (M+6), and valine (M+5) labeled. Data shown are mean ± S.D. (N = 4). **(C)** Y- axis of the bar graph shows the fold change in pool size of labeled BCAAs remaining in the culture media relative to WT, from the same experiment described in B. Data shown are mean ± S.D. (N = 4)

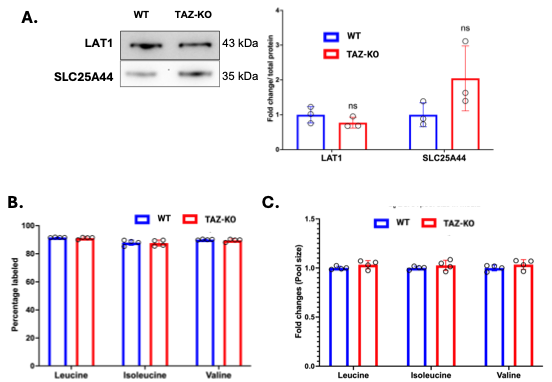

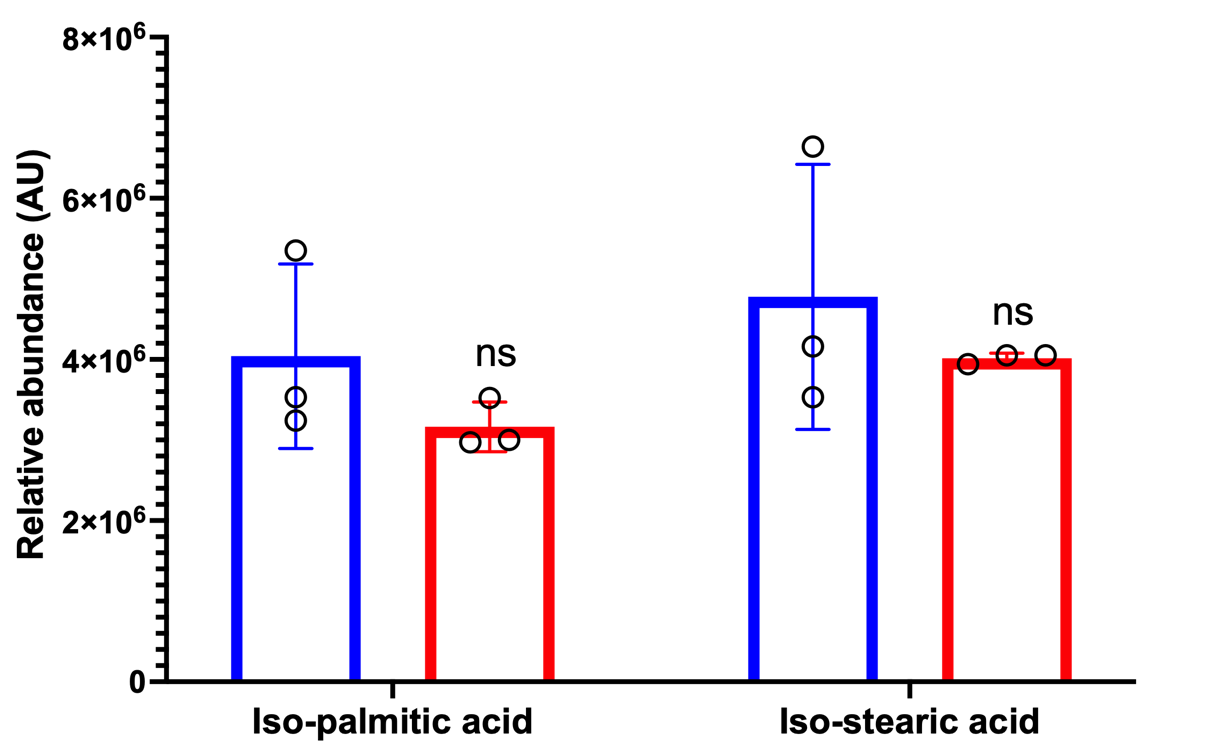

**Supporting info 2. ¹³C-BCAA incorporation into branched-chain fatty acids in WT and TAZ-KO cells.** WT and TAZ-KO cells were incubated with [U-¹³C]-leucine, [U-¹³C]-isoleucine, and [U-¹³C]-valine for 4 hours. Relative abundance of isopalmitic acid and isostearic acid was measured by LC-MS at the Lipidomics Core Facility at Wayne State University. x-axis shows individual BCFAs. y-axis represents relative abundance in arbitrary units (AU). Data are presented as mean ± S.D. (n=3). Statistical comparisons were made using unpaired, two-tailed Student's t-tests. ns = not significant.

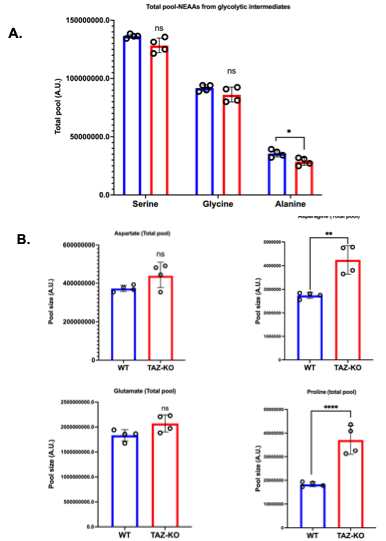

**Supplementary info 3: Total intracellular pools of NEAAs in WT and TAZ-KO C2C12 myoblasts.** Total pool sizes were determined by LC-MS in WT and TAZ-KO C2C12 myoblasts following a 4-h incubation with [U-¹³C₆]glucose and represent the sum of all isotopomers (M+0 through M+n). (A) Total pools of amino acids derived from glycolytic intermediates. (B) Total pools of amino acids derived from TCA cycle intermediates. The X-axis represents individual amino acids and the Y-axis represents total intracellular pool size normalized to cell number. Data shown are mean ± S.D. (n = 4). *, p < 0.01; **, p < 0.01; ***, p < 0.001.

**Supplementary info 4 (A-K). Total protein normalization images for western blot quantification.** Total protein was quantified from each sample using No-Stain labeling reagent (Invitrogen) and used for total protein normalization (TPN) of the corresponding western blots presented in the main figures. Supplementary figures A–K show the No-Stain total protein images corresponding to western blots.

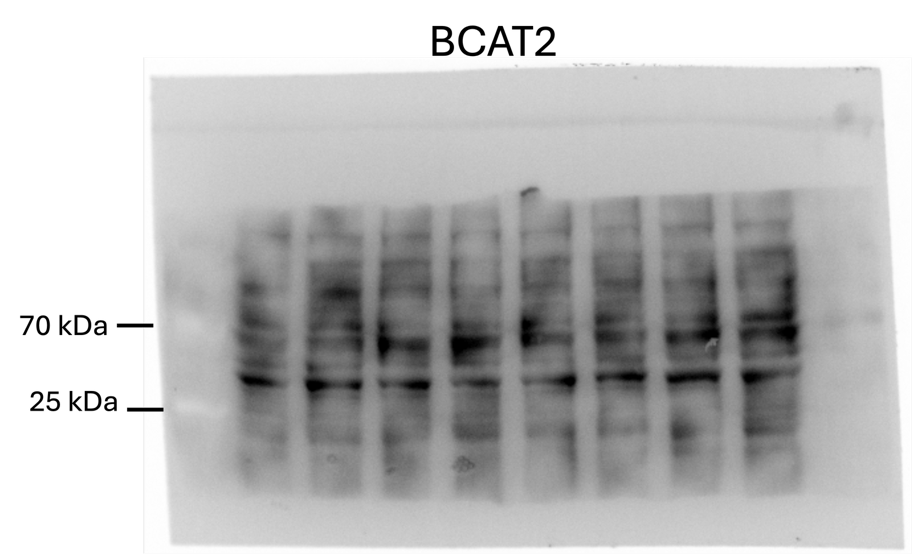

**4 A:**

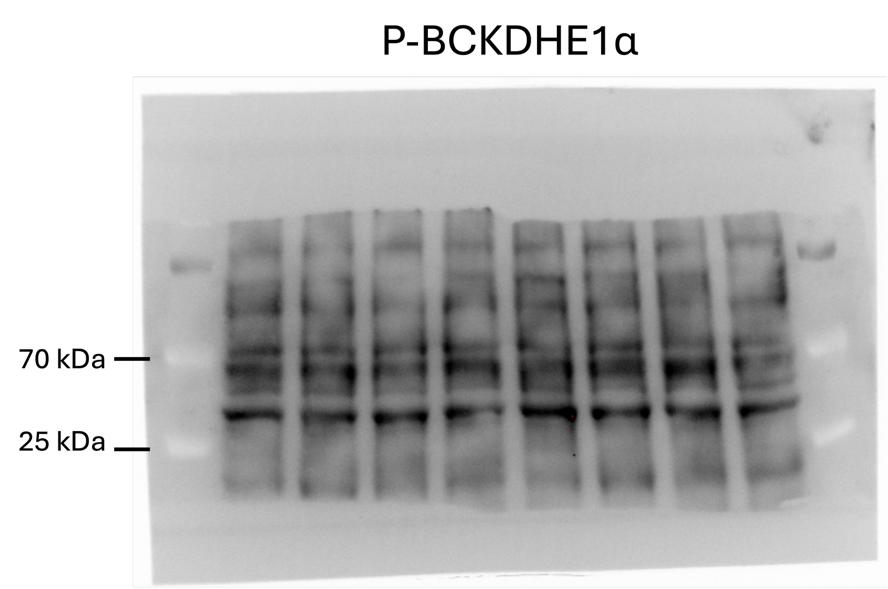

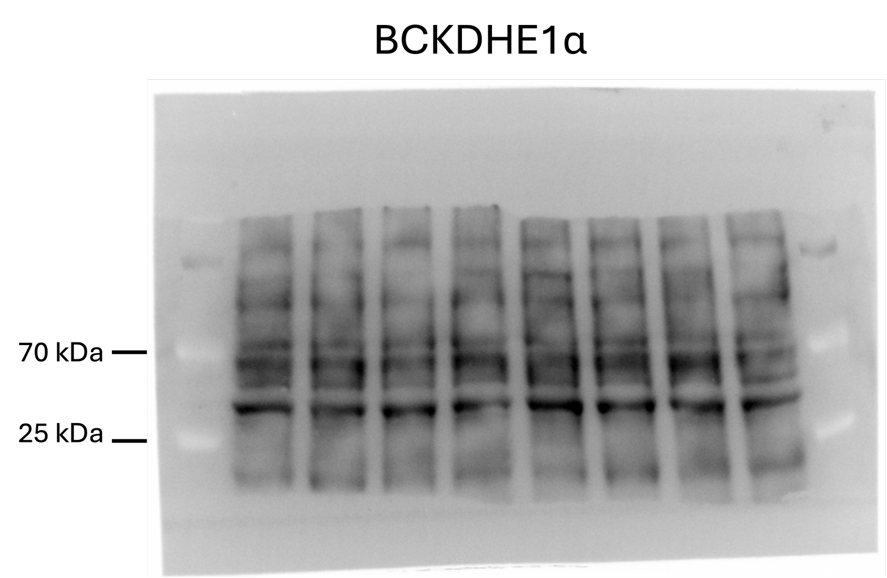
**4 B:**

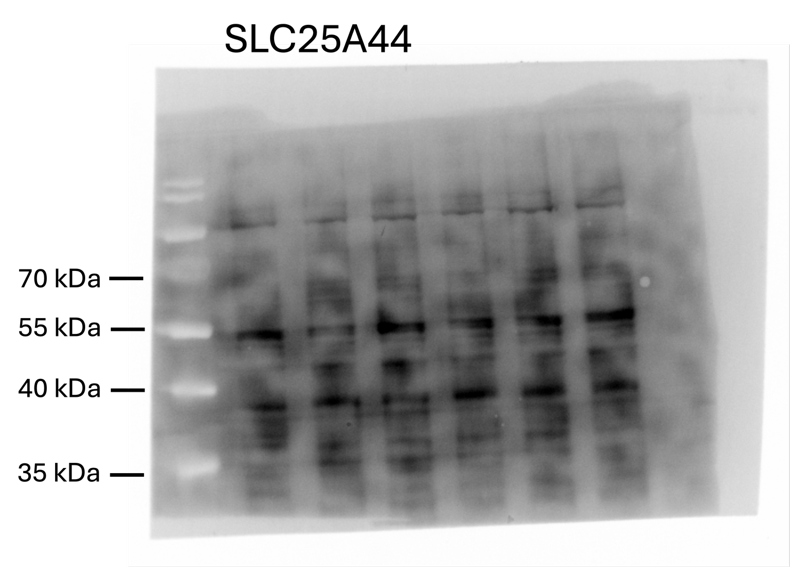
**4 C:**

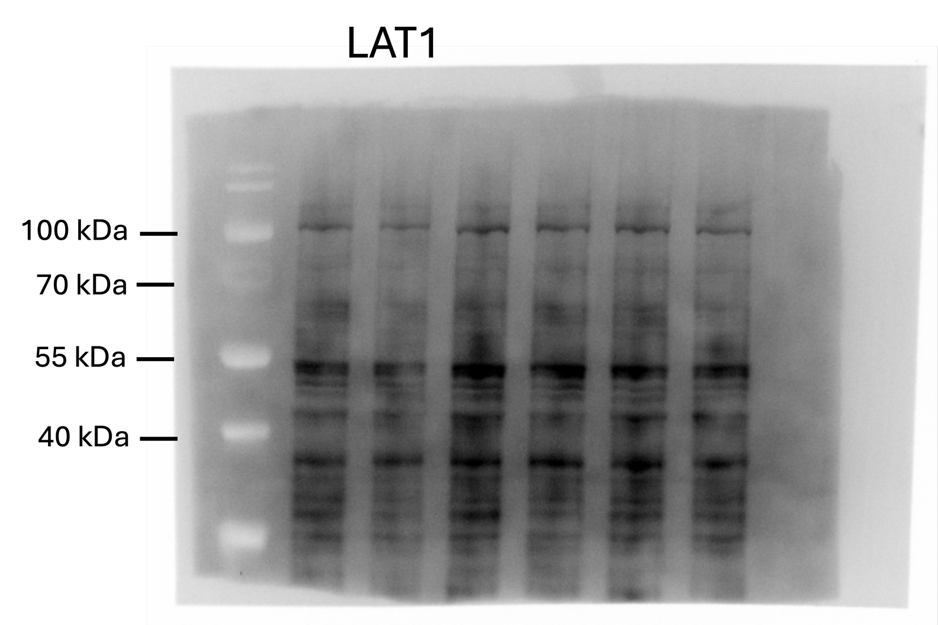

**4 D:**

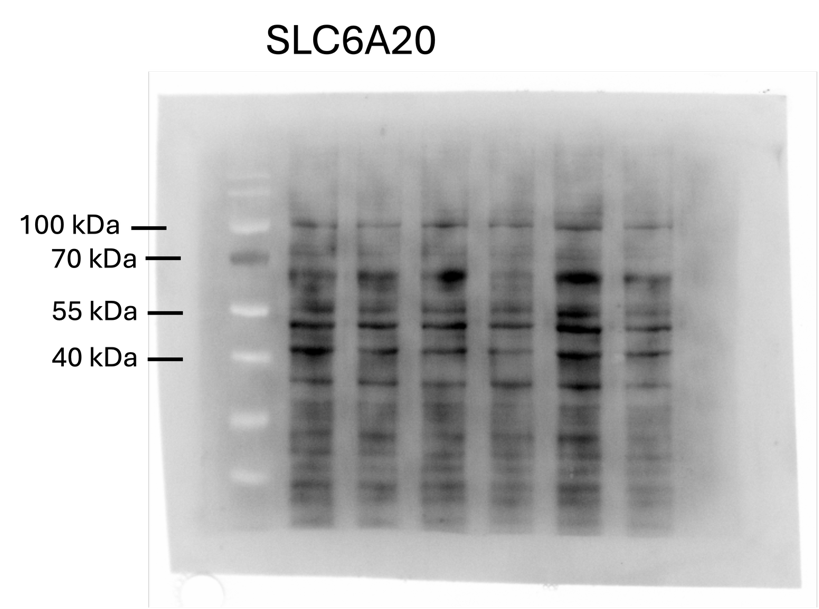
**4 E:**

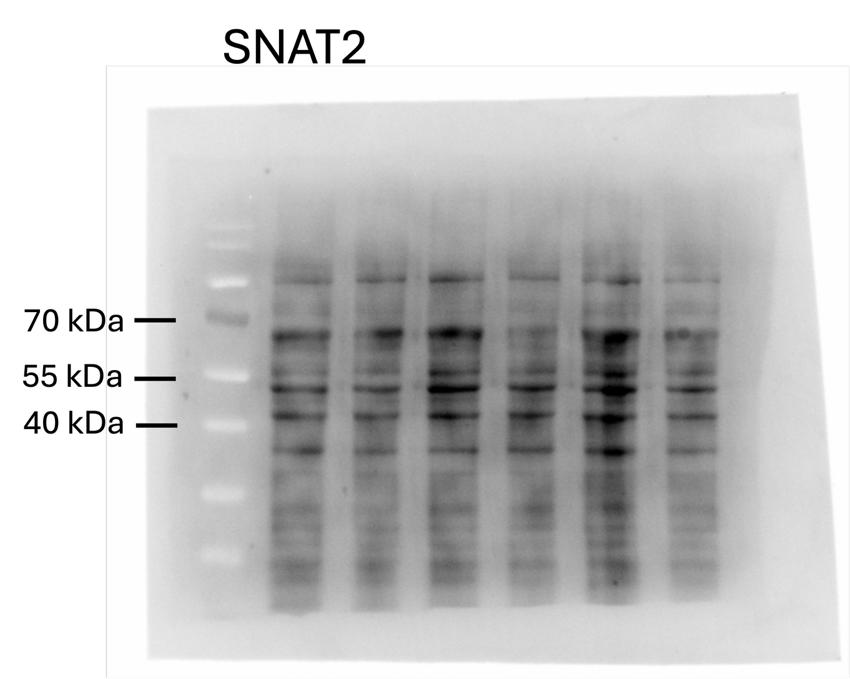

**4 F:**

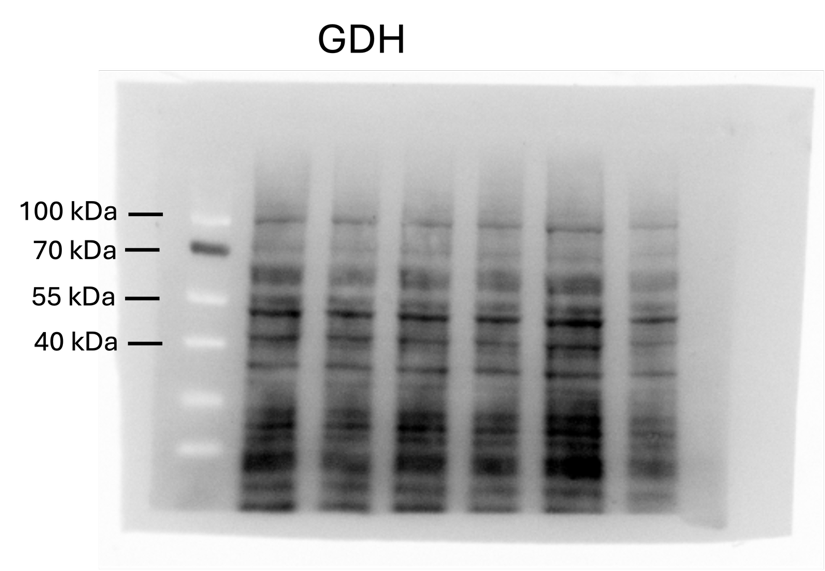
**4 G:**

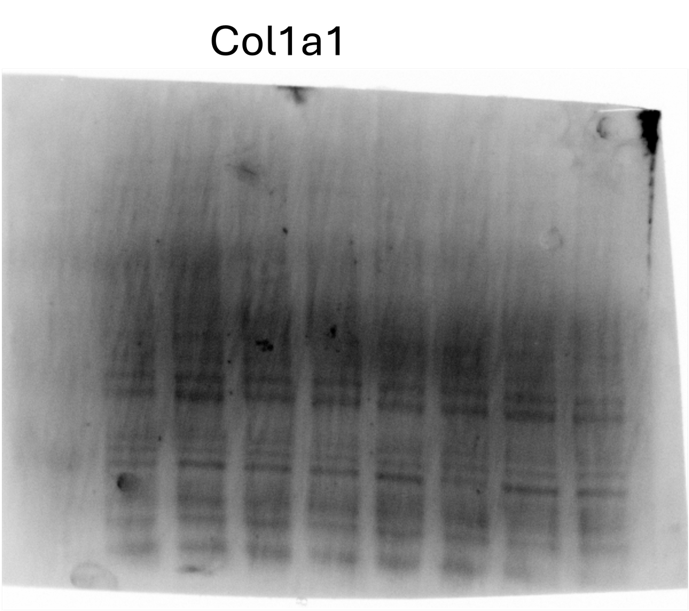

**4 H:**

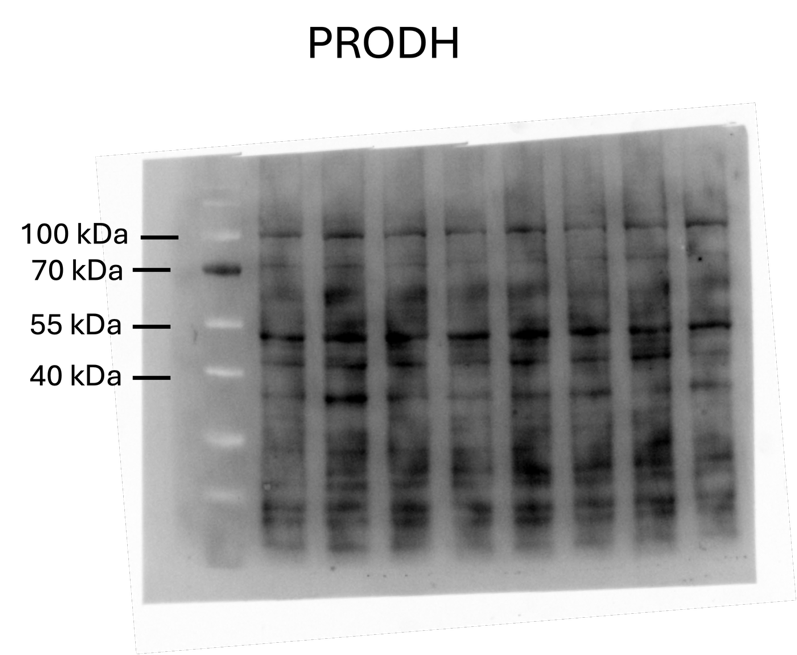
**4 I:**

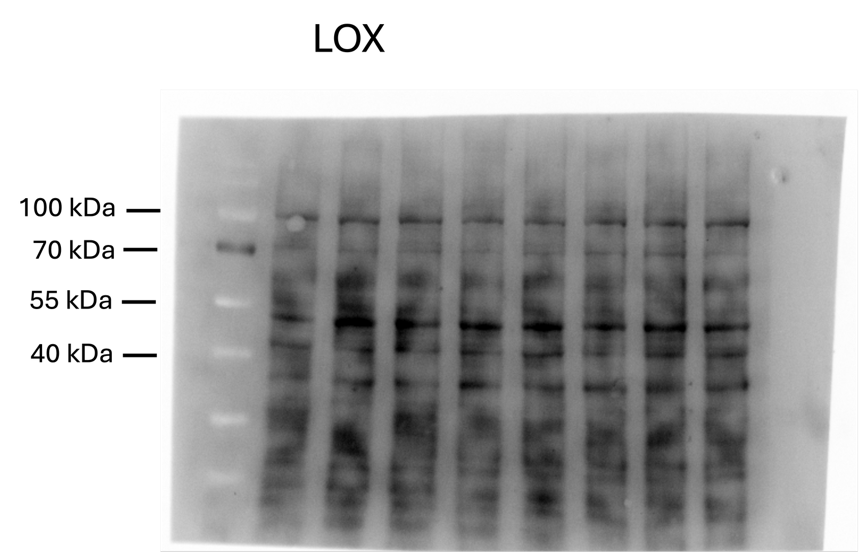

**4 J:**

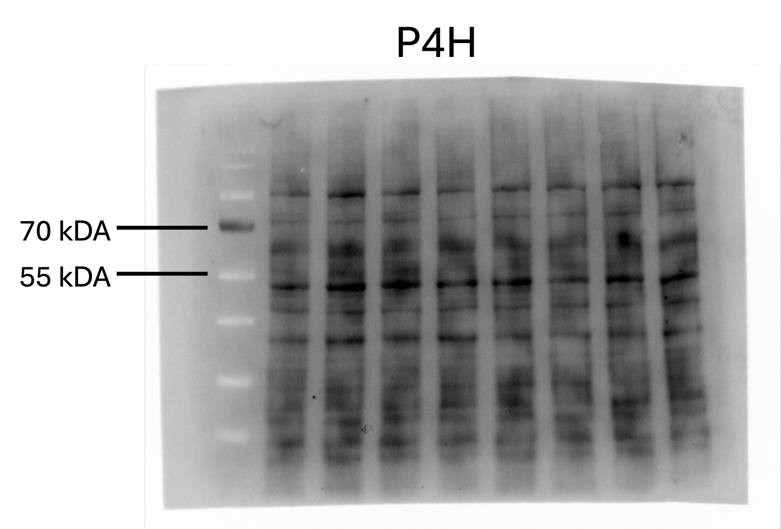

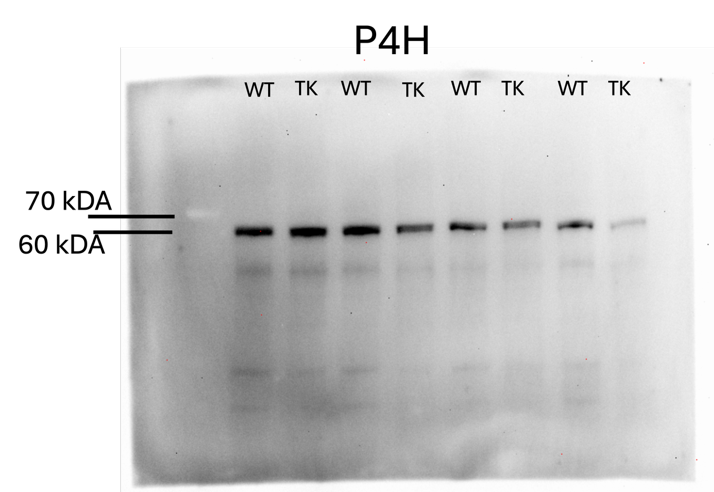
**4 K:**

| Table S1: Panel of compounds included for targeted MS2 triggering during mass spectrometry acquisition | | | | | | |
| --- | --- | --- | --- | --- | --- | --- |
| Compound | Formula | Adduct | m/z | z | RT Time (min) | Window (min) |
| 2,3-Dihydroxy-2-methylbutanoic acid | C5H10O4 | -H | 133.0506 | 1 | 10.4 | 2 |
| 2,3-Diketogulonic acid | C6H8O7 | -H | 191.0197 | 1 | 9.4 | 2 |
| 2-Hydroxybutyrate | C4H8O3 | -H | 103.0401 | 1 | 10.83 | 2 |
| 2-Methylcitrate | C7H10O7 | -H | 205.0354 | 1 | 15.8 | 2 |
| 3-D-Hydroxybutyrate | C4H8O3 | -H | 103.0401 | 1 | 9.19 | 2 |
| 3-Hydroxyisobutyrate | C4H8O3 | -H | 103.0401 | 1 | 9.1 | 2 |
| 3-Methyl-2-oxovaleric acid | C6H10O3 | -H | 129.0557 | 1 | 16.56 | 2 |
| 3-Phosphoglyceric acid | C3H7O7P | -H | 184.9857 | 1 | 15.3 | 2 |
| 4-Hydroxybutyrate | C4H8O3 | -H | 103.0401 | 1 | 10.3 | 2 |
| 4-Hydroxyproline | C5H9NO3 | -H | 130.051 | 1 | 1.44 | 2 |
| 5-Aminolevulinic acid | C5H9NO3 | -H | 130.051 | 1 | 1.42 | 2 |
| 5-Hydroxyindoleacetic acid | C10H9NO3 | -H | 190.051 | 1 | 13.58 | 2 |
| 5-L-Glutamyl-taurine | C7H14N2O6S | -H | 253.05 | 1 | 6.8 | 2 |
| 5'-Methylthioadenosine (MTA) | C11H15N5O3S | -H | 296.0823 | 1 | 12.66 | 2 |
| Acetyl-CoA | C23H38N7O17P3S | -H | 808.1185 | 1 | 17.88 | 2 |
| Adenosine 3-phosphate 5-phosphosulfate | C10H15N5O13P2S | -H | 505.979 | 1 | 17.1 | 2 |
| ADP-Ribose | C15H23N5O14P2 | -H | 558.0644 | 1 | 15.02 | 2 |
| alpha-Ketoglutarate | C5H6O5 | -H | 145.0142 | 1 | 14.74 | 2 |
| Asp-Asp | C8H12N2O7 | -H | 247.0572 | 1 | 14.02 | 2 |
| Asp-Glu | C9H14N2O7 | -H | 261.0728 | 1 | 13.9 | 2 |
| Asp-Gly | C6H10N2O5 | -H | 189.0517 | 1 | 5.4 | 2 |
| CDP-ethanolamine | C11H20N4O11P2 | -H | 445.0531 | 1 | 7.1 | 2 |
| CMP-n-acetylneuraminic acid | C20H31N4O16P | -H | 613.14 | 1 | 14.3 | 2 |
| D-2-Aminobutyrate | C4H9NO2 | -H | 102.0561 | 1 | 1.51 | 2 |
| Fructose 1,6-BP | C6H14O12P2 | -H | 338.9888 | 1 | 15.9 | 2 |
| gamma-Glu-Cys | C8H14N2O5S | -H | 249.0551 | 1 | 9 | 2 |
| gamma-Glu-Thr | C9H16N2O6 | -H | 247.0936 | 1 | 7.7 | 2 |
| gamma-Glu-Val | C10H18N2O5 | -H | 245.1143 | 1 | 13.4 | 2 |
| GDP-Mannose | C16H25N5O16P2 | -H | 604.0699 | 1 | 14.35 | 2 |
| Gln-Glu | C10H17N3O6 | -H | 274.1045 | 1 | 7.3 | 2 |
| Glu-Ala | C8H14N2O5 | -H | 217.083 | 1 | 9.3 | 2 |
| Glu-Gly | C7H12N2O5 | -H | 203.0673 | 1 | 6.45 | 2 |
| Glu-Ile | C11H20N2O5 | -H | 259.1299 | 1 | 14 | 2 |
| Glycerol 3-phosphate | C3H9O6P | -H | 171.0064 | 1 | 10.1 | 2 |
| Glycerol 3-phosphate | C3H9O6P | -H | 171.0064 | 1 | 10.3 | 2 |
| N-Acetyl-1-aspartylglutamic acid | C11H16N2O8 | -H | 303.0834 | 1 | 16.5 | 2 |
| N-Acetyl-D-hexosamine phosphate A | C8H16NO9P | -H | 300.049 | 1 | 10.4 | 2 |
| N-Acetyl-D-hexosamine phosphate B | C8H16NO9P | -H | 300.049 | 1 | 10.7 | 2 |
| N-Acetylglutamine | C7H12N2O4 | -H | 187.0724 | 1 | 8.7 | 2 |
| N-Acetyl-L-alanine | C5H9NO3 | -H | 130.051 | 1 | 9.84 | 2 |
| N-Acetyl-L-aspartic acid | C6H9NO5 | -H | 174.0408 | 1 | 14.43 | 2 |
| N-Acetyl-L-glutamic acid | C7H11NO5 | -H | 188.0564 | 1 | 14.53 | 2 |
| N-Acetyl-L-methionine | C7H13NO3S | -H | 190.0543 | 1 | 14.5 | 2 |
| N-Acetyl-L-phenylalanine | C11H13NO3 | -H | 206.0823 | 1 | 16.81 | 2 |
| N-Acetylneuraminic acid | C11H19NO9 | -H | 308.0987 | 1 | 8.1 | 2 |
| N-Acetylserine | C5H9NO4 | -H | 146.0459 | 1 | 7.9 | 2 |
| N-Acetyltryptophan | C13H14N2O3 | -H | 245.0932 | 1 | 16.6 | 2 |
| N-Glycolylneuraminic acid | C11H19NO10 | -H | 324.0936 | 1 | 7.8 | 2 |
| O-Acetylserine | C5H9NO4 | -H | 146.0459 | 1 | 7.97 | 2 |
| O-Phosphoethanolamine | C2H8NO4P | -H | 140.0118 | 1 | 1.96 | 2 |
| Pyridoxal 5'-phosphate | C8H10NO6P | -H | 246.0173 | 1 | 16.1 | 2 |
| Sedoheptulose 7-phosphate | C7H15O10P | -H | 289.033 | 1 | 10 | 2 |
| S-Lactoylglutathione | C13H21N3O8S | -H | 378.0977 | 1 | 15.7 | 2 |
| UDP-alpha-D-glucuronic acid | C15H22N2O18P2 | -H | 579.027 | 1 | 16.7 | 2 |
| UDP-hexose | C15H24N2O17P2 | -H | 565.0477 | 1 | 14.34 | 2 |
| UDP-N-acetylhexosamine | C17H27N3O17P2 | -H | 606.0743 | 1 | 14.44 | 2 |
| Adenine | C5H5N5 | -H | 134.0472 | 1 | 4.3 | 2 |
| Adenosine | C10H13N5O4 | -H | 266.0895 | 1 | 8.03 | 2 |
| ADP | C10H15N5O10P2 | -H | 426.0221 | 1 | 15.7 | 2 |
| AICAR | C9H15N4O8P | -H | 337.0555 | 1 | 14.1 | 2 |
| Alanine | C3H7NO2 | -H | 88.0404 | 1 | 1.38 | 2 |
| AMP | C10H14N5O7P | -H | 346.0558 | 1 | 13.3 | 2 |
| Arachidonic acid | C20H32O2 | -H | 303.233 | 1 | 23.5 | 1 |
| Ascorbate | C6H8O6 | -H | 175.0248 | 1 | 7.4 | 2 |
| Asparagine | C4H8N2O3 | -H | 131.0462 | 1 | 1.36 | 2 |
| Aspartate | C4H7NO4 | -H | 132.0302 | 1 | 6.1 | 2 |
| ATP | C10H16N5O13P3 | -H | 505.9885 | 1 | 17.1 | 2 |
| Bilirubin | C33H36N4O6 | -H | 583.2562 | 1 | 22.42 | 2 |
| Bisphosphoglycerate | C3H8O10P2 | -H | 264.952 | 1 | 17.1 | 2 |
| cAMP | C10H12N5O6P | -H | 328.0452 | 1 | 14 | 2 |
| Carglumic acid | C6H10N2O5 | -H | 189.0517 | 1 | 14.05 | 2 |
| CDP | C9H15N3O11P2 | -H | 402.0109 | 1 | 15.1 | 2 |
| Cellobiose | C12H22O11 | -H | 341.1089 | 1 | 1.49 | 2 |
| Citraconic acid | C5H6O4 | -H | 129.0193 | 1 | 15.8 | 2 |
| Citrate | C6H8O7 | -H | 191.0197 | 1 | 16 | 2 |
| Citrulline | C6H13N3O3 | -H | 174.0884 | 1 | 1.42 | 2 |
| CMP | C9H14N3O8P | -H | 322.0446 | 1 | 11.4 | 2 |
| Coenzyme A | C21H36N7O16P3S | -H | 766.1079 | 1 | 17.7 | 2 |
| Creatine | C4H9N3O2 | -H | 130.0622 | 1 | 1.43 | 2 |
| Creatinine | C4H7N3O | -H | 112.0516 | 1 | 1.55 | 2 |
| CTP | C9H16N3O14P3 | -H | 481.9772 | 1 | 16.8 | 2 |
| Cystathionine | C7H14N2O4S | -H | 221.0602 | 1 | 1.4 | 2 |
| Cytidine | C9H13N3O5 | -H | 242.0782 | 1 | 2.21 | 2 |
| Cytosine | C4H5N3O | -H | 110.036 | 1 | 1.57 | 2 |
| dCMP | C9H14N3O7P | -H | 306.0497 | 1 | 11.9 | 2 |
| Dehydroascorbic acid | C6H6O6 | -H | 173.0092 | 1 | 1.85 | 2 |
| Deoxyuridine | C9H12N2O5 | -H | 227.0673 | 1 | 5.44 | 2 |
| DHAP | C3H7O6P | -H | 168.9907 | 1 | 12.06 | 2 |
| Dihydroorotic acid | C5H6N2O4 | -H | 157.0255 | 1 | 8.19 | 2 |
| Dihydroxyacetone phosphate | C3H7O6P | -H | 168.9907 | 1 | 12.2 | 2 |
| Dimethylglycine | C4H9NO2 | -H | 102.0561 | 1 | 1.43 | 2 |
| Dopamine | C8H11NO2 | -H | 152.0717 | 1 | 1.08 | 2 |
| dTDP | C10H16N2O11P2 | -H | 401.0157 | 1 | 15.7 | 2 |
| dTMP | C10H15N2O8P | -H | 321.0493 | 1 | 13 | 2 |
| dTTP | C10H17N2O14P3 | -H | 480.982 | 1 | 17.1 | 2 |
| FAD | C27H33N9O15P2 | -H | 784.1499 | 1 | 16.9 | 2 |
| Farnesyl pyrophosphate | C15H28O7P2 | -H | 381.1237 | 1 | 21.2 | 2 |
| FMN | C17H21N4O9P | -H | 455.0973 | 1 | 16.11 | 2 |
| Folic acid | C19H19N7O6 | -H | 440.1324 | 1 | 15.61 | 2 |
| Fumaric acid | C4H4O4 | -H | 115.0037 | 1 | 14.9 | 2 |
| GABA | C4H9NO2 | -H | 102.0561 | 1 | 1.41 | 2 |
| Galactonic acid | C6H12O7 | -H | 195.051 | 1 | 7.2 | 2 |
| GDP | C10H15N5O11P2 | -H | 442.0171 | 1 | 15.3 | 2 |
| Gluconic acid | C6H12O7 | -H | 195.051 | 1 | 7.3 | 2 |
| Glucose | C6H12O6 | -H | 179.0561 | 1 | 1.44 | 2 |
| Glutaconic acid | C5H6O4 | -H | 129.0193 | 1 | 16 | 2 |
| Glutamate | C5H9NO4 | -H | 146.0459 | 1 | 5.66 | 2 |
| Glutamine | C5H10N2O3 | -H | 145.0619 | 1 | 1.37 | 2 |
| Glutathione (oxidized) | C20H32N6O12S2 | -H | 611.1447 | 1 | 13.95 | 2 |
| Glutathione (reduced) | C10H17N3O6S | -H | 306.0765 | 1 | 9.46 | 2 |
| Glycine | C2H5NO2 | -H | 74.0248 | 1 | 1.41 | 2 |
| GMP | C10H14N5O8P | -H | 362.0507 | 1 | 12.3 | 2 |
| GTP | C10H16N5O14P3 | -H | 521.9834 | 1 | 16.9 | 2 |
| Hexose phosphate A | C6H13O9P | -H | 259.0224 | 1 | 9.4 | 2 |
| Hexose phosphate B | C6H13O9P | -H | 259.0224 | 1 | 9.6 | 2 |
| Hexose phosphate C | C6H13O9P | -H | 259.0224 | 1 | 10 | 2 |
| Hexose phosphate D | C6H13O9P | -H | 259.0224 | 1 | 10.3 | 2 |
| Hexose phosphate E | C6H13O9P | -H | 259.0224 | 1 | 10.4 | 2 |
| Hexose phosphate F | C6H13O9P | -H | 259.0224 | 1 | 11.6 | 2 |
| Hexose phosphate G | C6H13O9P | -H | 259.0224 | 1 | 12.1 | 2 |
| Hexose phosphate H | C6H13O9P | -H | 259.0224 | 1 | 12.5 | 2 |
| Hexose phosphate I | C6H13O9P | -H | 259.0224 | 1 | 13.2 | 2 |
| Hexose phosphate J | C6H13O9P | -H | 259.0224 | 1 | 15.1 | 2 |
| Histidine | C6H9N3O2 | -H | 154.0622 | 1 | 1.13 | 2 |
| Hydroxyphenyllactic acid | C9H10O4 | -H | 181.0506 | 1 | 13.4 | 2 |
| Hypoxanthine | C5H4N4O | -H | 135.0312 | 1 | 1.96 | 2 |
| IMP | C10H13N4O8P | -H | 347.0398 | 1 | 12.3 | 2 |
| Indolelactic acid | C11H11NO3 | -H | 204.0666 | 1 | 16.5 | 2 |
| Indoxyl sulfate | C8H7NO4S | -H | 212.0023 | 1 | 16.66 | 2 |
| Inosine | C10H12N4O5 | -H | 267.0735 | 1 | 6.4 | 2 |
| Isoleucine | C6H13NO2 | -H | 130.0874 | 1 | 2.92 | 2 |
| Itaconate | C5H6O4 | -H | 129.0193 | 1 | 14.11 | 2 |
| Ketoisovalerate | C5H8O3 | -H | 115.0401 | 1 | 14.6 | 2 |
| Ketoleucine | C6H10O3 | -H | 129.0557 | 1 | 16.77 | 2 |
| Lactate | C3H6O3 | -H | 89.0244 | 1 | 8.31 | 2 |
| Leucine | C6H13NO2 | -H | 130.0874 | 1 | 3.21 | 2 |
| Malate | C4H6O5 | -H | 133.0142 | 1 | 14.5 | 2 |
| Malonic acid | C3H4O4 | -H | 103.0037 | 1 | 13.9 | 2 |
| Methionine | C5H11NO2S | -H | 148.0438 | 1 | 1.96 | 2 |
| NAD+ | C21H27N7O14P2 | -H | 662.1018 | 1 | 10.4 | 2 |
| NADH | C21H29N7O14P2 | -H | 664.1175 | 1 | 15.66 | 2 |
| NADP+ | C21H28N7O17P3 | -H | 742.0682 | 1 | 15.7 | 2 |
| NADPH | C21H30N7O17P3 | -H | 744.0838 | 1 | 17.2 | 2 |
| Ophthalmate | C11H19N3O6 | -H | 288.1201 | 1 | 9.6 | 2 |
| Orotic acid | C5H4N2O4 | -H | 155.0098 | 1 | 10.1 | 2 |
| Orotidine | C10H12N2O8 | -H | 287.0521 | 1 | 9.1 | 2 |
| Pantothenic acid | C9H17NO5 | -H | 218.1034 | 1 | 12.5 | 2 |
| Pentose phosphate A | C5H11O8P | -H | 229.0119 | 1 | 9.7 | 2 |
| Pentose phosphate B | C5H11O8P | -H | 229.0119 | 1 | 11 | 2 |
| Pentose phosphate C | C5H11O8P | -H | 229.0119 | 1 | 11.1 | 2 |
| Pentose phosphate D | C5H11O8P | -H | 229.0119 | 1 | 11.7 | 2 |
| Pentose phosphate E | C5H11O8P | -H | 229.0119 | 1 | 12.3 | 2 |
| Phenylalanine | C9H11NO2 | -H | 164.0717 | 1 | 6.89 | 2 |
| Phenyllactic acid | C9H10O3 | -H | 165.0557 | 1 | 16.6 | 2 |
| Phosphocreatine | C4H10N3O5P | -H | 210.0285 | 1 | 14.1 | 2 |
| Phosphoenolpyruvate | C3H5O6P | -H | 166.9751 | 1 | 15.9 | 2 |
| Phosphoribosyl pyrophosphate | C5H13O14P3 | -H | 388.9445 | 1 | 17 | 2 |
| Phosphoserine | C3H8NO6P | -H | 184.0016 | 1 | 11.96 | 2 |
| Proline | C5H9NO2 | -H | 114.0561 | 1 | 1.44 | 2 |
| Pyridoxal | C8H9NO3 | -H | 166.051 | 1 | 3.1 | 2 |
| Pyridoxine | C8H11NO3 | -H | 168.0666 | 1 | 3.39 | 2 |
| Pyroglutamic acid | C5H7NO3 | -H | 128.0353 | 1 | 8.92 | 2 |
| Pyruvate | C3H4O3 | -H | 87.0088 | 1 | 10.4 | 2 |
| Riboflavin | C17H20N4O6 | -H | 375.131 | 1 | 13.35 | 2 |
| SAH | C14H20N6O5S | -H | 383.1143 | 1 | 7 | 2 |
| Serine | C3H7NO3 | -H | 104.0353 | 1 | 1.36 | 2 |
| Succinate | C4H6O4 | -H | 117.0193 | 1 | 14.1 | 2 |
| Taurine | C2H7NO3S | -H | 124.0074 | 1 | 1.4 | 2 |
| Threonic acid | C4H8O5 | -H | 135.0299 | 1 | 6.98 | 2 |
| Threonine | C4H9NO3 | -H | 118.051 | 1 | 1.39 | 2 |
| Thymidine | C10H14N2O5 | -H | 241.083 | 1 | 6.6 | 2 |
| Tricarballylic acid | C6H8O6 | -H | 175.0248 | 1 | 15.21 | 2 |
| Tryptophan | C11H12N2O2 | -H | 203.0826 | 1 | 9.03 | 2 |
| Tyrosine | C9H11NO3 | -H | 180.0666 | 1 | 3.27 | 2 |
| UDP | C9H14N2O12P2 | -H | 402.9949 | 1 | 15.3 | 2 |
| UMP | C9H13N2O9P | -H | 323.0286 | 1 | 12.08 | 2 |
| Uracil | C4H4N2O2 | -H | 111.02 | 1 | 2.16 | 2 |
| Ureidosuccinic acid | C5H8N2O5 | -H | 175.036 | 1 | 14.29 | 2 |
| Uric acid | C5H4N4O3 | -H | 167.0211 | 1 | 6.98 | 2 |
| Uridine | C9H12N2O6 | -H | 243.0623 | 1 | 3.85 | 2 |
| UTP | C9H15N2O15P3 | -H | 482.9613 | 1 | 16.9 | 2 |
| Valine | C5H11NO2 | -H | 116.0717 | 1 | 1.6 | 2 |
| Xanthine | C5H4N4O2 | -H | 151.0261 | 1 | 2.18 | 2 |
| Xanthosine | C10H12N4O6 | -H | 283.0684 | 1 | 9.85 | 2 |

| **Table S2: Formulation for Dulbecco's Modified Eagle's Medium (DMEM) without BCAAs** | |
| --- | --- |
| **Component** | **Concentration (g/liter)** |
| **Inorganic Salts** | |
| CaCl₂ (anhydrous) | 0.20000 |
| Fe(NO₃)₃·9H₂O | 0.00010 |
| MgSO₄ (anhydrous) | 0.09770 |
| KCl | 0.40000 |
| NaHCO₃ | 1.50000 |
| NaCl | 6.40000 |
| NaH₂PO₄·H₂O | 0.12500 |
| **Amino Acids** | |
| L-Arginine·HCl | 0.08400 |
| L-Cystine·2HCl | 0.06260 |
| L-Glutamine | 0.58400 |
| L-Glycine | 0.03000 |
| L-Histidine·HCl·H₂O | 0.04200 |
| ***L-Isoleucine*** | - |
| ***L-Leucine*** | - |
| L-Lysine·HCl | 0.14600 |
| L-Methionine | 0.03000 |
| L-Phenylalanine | 0.06600 |
| L-Serine | 0.04200 |
| L-Threonine | 0.09500 |
| L-Tryptophan | 0.01600 |
| L-Tyrosine·2Na·2H₂O | 0.10379 |
| ***L-Valine*** | - |
| **Vitamins** | |
| Choline Chloride | 0.00400 |
| Folic Acid | 0.00400 |
| myo-Inositol | 0.00720 |
| Nicotinamide | 0.00400 |
| D-Pantothenic Acid (hemicalcium) | 0.00400 |
| Pyridoxine·HCl | 0.00400 |
| Riboflavin | 0.00040 |
| Thiamine·HCl | 0.00400 |
| **Other** | |
| D-Glucose | 4.50000 |
| Phenol Red, Sodium Salt | 0.01500 |
| Sodium Pyruvate | 0.11000 |
